# SMN deficiency induces an early non-atrophic myopathy with alterations in the contractile and excitatory coupling machinery of skeletal myofibers in the SMNΔ7 mouse model of spinal muscular atrophy

**DOI:** 10.1101/2024.09.11.612470

**Authors:** María T. Berciano, Alaó Gatius, Alba Puente-Bedia, Alexis Rufino-Gómez, Olga Tarabal, José C. Rodríguez-Rey, Jordi Calderó, Miguel Lafarga, Olga Tapia

## Abstract

Spinal muscular atrophy (SMA) is caused by a deficiency of the ubiquitously expressed survival motor neuron (SMN) protein. The main pathological hallmark of SMA is the degeneration of lower motor neurons (MNs) with subsequent denervation and atrophy of skeletal muscle. However, increasing evidence indicates that low SMN levels not only are detrimental to the central nervous system but also directly affect other peripheral tissues and organs, including skeletal muscle. To better understand the potential primary impact of SMN deficiency in muscle, we explored the cellular, ultrastructural, and molecular basis of SMA myopathy in the SMNΔ7 mouse model of severe SMA at an early postnatal period (P0-7) prior to muscle denervation and MN loss (preneurodegenerative [PND] stage). This period contrasts with the neurodegenerative (ND) stage (P8-14), in which MN loss and muscle atrophy occur. At the PND stage, we found that SMNΔ7 mice displayed early signs of motor dysfunction with overt myofiber alterations in the absence of atrophy. Focal and segmental lesions in the myofibrillar contractile apparatus were noticed in myofibers. These lesions were observed in association with specific myonuclear domains and included abnormal accumulations of actin-thin myofilaments, sarcomere disruption, and the formation of minisarcomeres. The sarcoplasmic reticulum and triads also exhibited ultrastructural alterations, suggesting decoupling during the excitation-contraction process. Finally, changes in intermyofibrillar mitochondrial organization and dynamics, indicative of mitochondrial biogenesis overactivation, were also found. Overall, our results demonstrated that SMN deficiency induces early and MN loss-independent alterations in myofibers that essentially contribute to SMA myopathy. This strongly supports the view of an intrinsic alteration of skeletal muscle in SMA, suggesting that this peripheral tissue is a key therapeutic target for the disease.

**Graphical summary:** Schematic representation of the main cellular and ultrastructural changes occurring in *tibialis anterior* (TA) myofibers of the SMNΔ7 mouse model of severe SMA during the pre-neurodegenerative stage (PND, P0-P7). See that the PND stage is characterized by the absence of MN loss and muscular atrophy but, SMN deficient myofibers display multiple alterations. Observe the early impact of low SMN levels on sarcoplasmic reticulum and triads, myofibril contractile cytoskeleton, and intermyofibrillar mitochondria. Notice that all alterations are associated with a specific myonuclear domain. Changes in mRNA levels of different genes involved in myogenesis and mitochondrial biogenesis are also shown compared to age-matched WT animals. Figure created with BioRender.com.

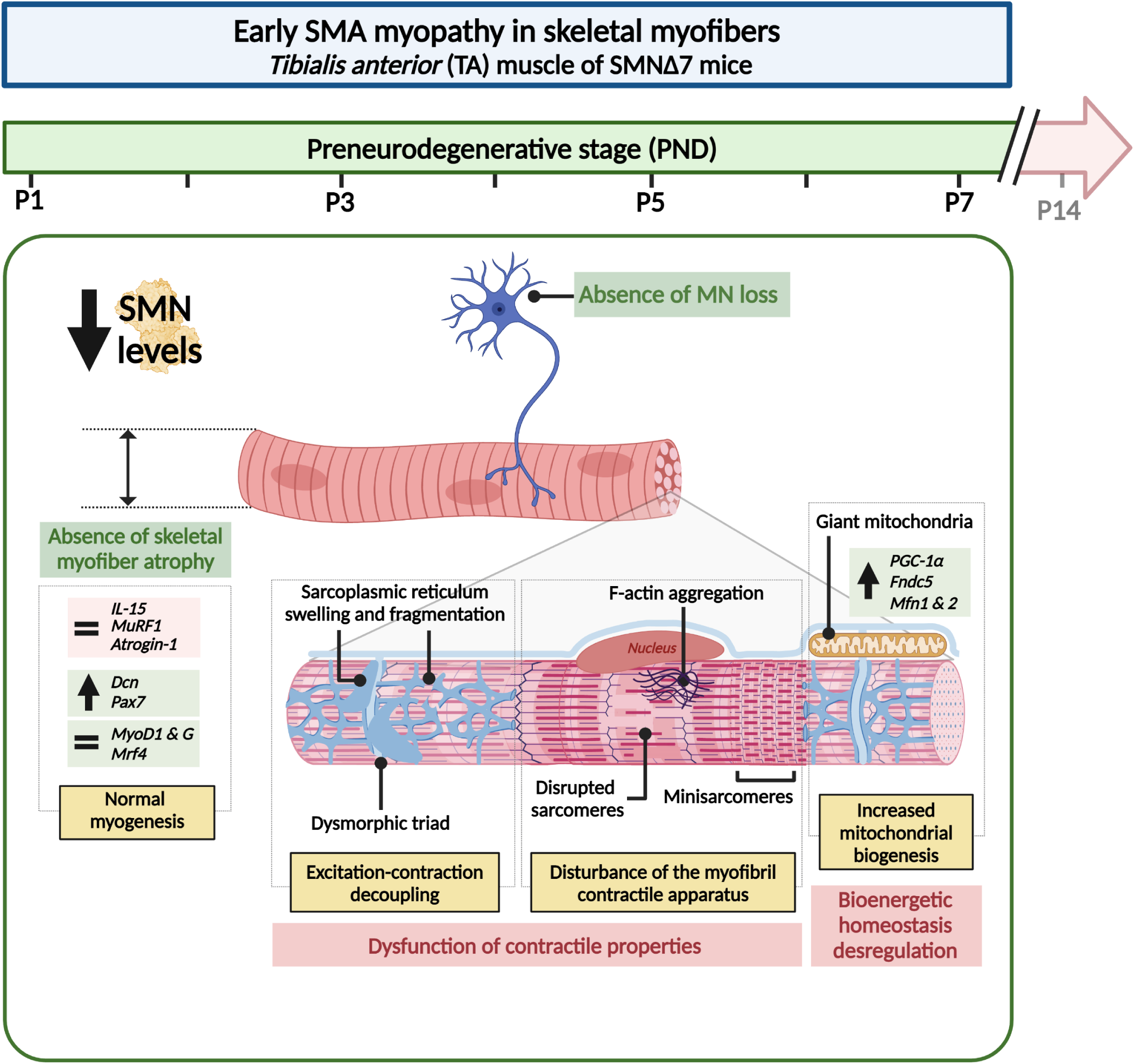

## INTRODUCTION

Spinal muscular atrophy (SMA) is an autosomal recessive neurodegenerative disorder characterized by progressive weakness and atrophy of skeletal muscles resulting from the loss or dysfunction of spinal alpha-motor neurons (MNs) (Lunn and Wang, 2008; Prior, 2010; Talbot and Tizzano, 2017). SMA is caused by reduced levels of the survival motor neuron (SMN) protein due to deletion of or inactivating mutations in the *survival motor neuron 1* (*SMN1*) gene. The human genome harbors an almost identical copy of *SMN1,* referred to as *SMN2,* which differs from *SMN1* in the substitution of a single base pair in the sequence of exon 7 (c.840 C to T). This change results in exon 7 skipping in approximately 70%-80% of *SMN2* transcripts. The translation of these transcripts results in a truncated and nonfunctional protein (SMNΔ7) that is rapidly degraded by the proteasome (Lefebvre et al., 1995; Lorson et al., 1999; Monani et al., 1999). The copy number of *SMN2* varies among SMA patients, making this gene an important modifier of disease severity.

Among other functions (Chaytow et al., 2018; Singh et al., 2017), the SMN protein is essential for the biogenesis of small nuclear ribonucleoproteins (snRNPs), which are involved in the splicing of pre-mRNAs (Carissimi et al., 2006; Gabanella et al., 2007; Paushkin et al., 2002). Low levels of SMN cause defects in pre-mRNAs splicing not only in the CNS but also in other tissues and organs, such as the skeletal muscle, heart, autonomic and enteric nervous systems, and lymphatic organs. This indicates that SMA is a general splicing disease with multiorgan alterations (Boyer et al., 2013a; Hamilton and Gillingwater, 2013; Motyl et al., 2020; Nash et al., 2016; Singh et al., 2017; Zhang et al., 2008).

At present, there is growing evidence that SMN deficiency directly impacts muscle homeostasis (Avila et al., 2007; Boyer et al., 2013a; Chemello et al., 2023; Cifuentes-Diaz et al., 2001; Dachs et al., 2011; Deguise et al., 2016; Groen et al., 2018; Hayhurst et al., 2012; Lee et al., 2011; Murray et al., 2008; Mutsaers et al., 2011). In this regard, experiments in cultured myoblasts/myotubes derived from SMA patients and mouse models of this disease have shown that reduced levels of SMN affect myotube formation and maturation (Bricceno et al., 2014; Hellbach et al., 2018; Martínez-Hernández et al., 2009; Shafey et al., 2005). Moreover, studies in mouse and *Drosophila* skeletal myofibers have reported that SMN is a constitutive component of sarcomeres (Rajendra et al., 2007; Walker et al., 2008). More recently, we reported that SMN is specifically located in the M-line and I-band in human sarcomeres (Berciano et al., 2020a), suggesting that SMN contributes to the maintenance of sarcomere architecture.

Additional evidence of the intrinsic role of SMN in SMA myopathy comes from the observation that depletion of SMN in type I SMA patients induces sarcomeric disarray not only in atrophic denervated skeletal myofibers (hereafter referred to as myofibers) but also in innervated hypertrophied ones (Berciano et al., 2020a). Furthermore, in SMA mouse models, the restoration of SMN levels exclusively in the CNS with therapeutic antisense oligonucleotides only partially rescues motor function (Berciano et al., 2020b; Hua et al., 2011). However, peripheral SMN restoration in SMA mice notably compensates for its deficit in the spinal cord, improving motor function and mouse survival and enhancing the production of neurotrophic factors by peripheral tissues (Hua et al., 2015, 2011). Collectively, these data support that SMN deficiency contributes to SMA myopathy even in the absence of MN degeneration.

In the present study, we investigated the cellular and molecular basis of a potential primary myopathy induced by SMN depletion in the SMNΔ7 mouse model of SMA. We characterized a preneurodegenerative (PND) stage of SMA myopathy with early motor dysfunction and myofiber alterations in the absence of significant MN loss and muscular atrophy. We described the occurrence of ultrastructural changes in the contractile (sarcomeres) and excitation-contraction (sarcoplasmic reticulum and triads) machinery of myofibers. We also reported a relationship between the altered phenotype of SMA myofibers and changes in the expression levels of genes involved in myogenesis, mitochondrial dynamics and muscle atrophy. We propose that these early alterations in SMN-deficient myofibers are essential contributors to motor impairment in SMA and consider skeletal muscle a major therapeutic target, particularly at the initial stages of the disease.

## MATERIALS AND METHODS

### SMA animals

The *Smn*^+/−^*;SMN2*^+/+^*;SMN*Δ7^+/+^, heterozygous knockouts for *Smn,* were purchased from The Jackson Laboratory (Sacramento, CA, USA; stock number 005025), and crossed to generate SMNΔ7 (*Smn^−/−^*;*SMN2^+/+^;SMN*Δ*7^+/+^*) and WT (*Smn^+/+^;SMN2^+/+^;SMN*Δ*7^+/+^*) mice. For genotyping, DNA from mouse tail clip samples was analyzed using the Phire Tissue direct PCR mastermix kit (Thermo Fisher Scientific, Waltham, MA, USA) and the following primers: WT forward 5′-TCCAGCTCCGGGATATTGGGATTG-3′, SMNΔ7 reverse 5′-GGTAACGCCAGGGTTTTCC-3′ and WT reverse 5′ -TTTCTTCTGGCTGTGCCTTT-3′.

For motor behavior assessment, the righting reflex test was conducted on WT and SMNΔ7 mice at different postnatal ages as described in (Berciano et al., 2020b). This test assesses the motor ability for a mouse pup to be able to flip onto its feet from a supine position. Briefly, animals were placed face up on a bench pad, keeping them in this position for 3 s. After being released, the time to return to the prone position was recorded, giving a maximum time of 30 s for each trial. The test was repeated twice for each animal. For body weight analysis, WT and SMNΔ7 mice (at least 8 animals per group) were weighted at different postnatal ages.

Animal care and handling were performed in accordance with the Spanish legislation (Spanish Royal Decree 53/2013 BOE) and the guidelines of the European Commission for the Accommodation and Care of Laboratory Animals (revised in Appendix A of the Council Directive 2010/63/UE). The experimental plan was examined and approved by the Ethics Committee of the University of Cantabria and the Committee for Animal Care and Use of the University of Lleida.

### Immunofluorescence and confocal microscopy

Mice were fixed by perfusion with freshly prepared 3.7% paraformaldehyde (PFA) under deep anesthesia with pentobarbital (50 mg/kg). The spinal cords and tibialis anterior (TA) muscle were dissected and cryoprotected by immersion in 30% sucrose diluted in phosphate buffer saline (PBS: 137 mM NaCl, 2.7 mM KCl, 8 mM Na_2_HPO_4_, and 2 mM KH_2_PO_4_; pH 7.4). Tissue samples were embedded in Tissue-Tek OCT Compound (Sakura FineTek USA, Torrance, CA, USA, and frozen at −80°C until used.

For immunofluorescence, 8µm-thick transverse cryostat sections of spinal cord or TA muscle were mounted on SuperFrost slides, permeabilized with 0.5% Triton X-100 for 30 min, and counterstained with the following dyes: propidium iodide (PI, Sigma-Aldrich, Saint Louis, MO, USA) to label Nissl granules, Phalloidin-FITC (Sigma) to label actin filaments, MitoTracker (Invitrogen, Waltham, MA, USA) to label mitochondria or 4′,6-diamidino-2-phenylindole (DAPI, Sigma-Aldrich) to label nuclei. The slides were mounted with ProLong Anti-Fading Medium (Thermo Fisher Scientific). For image acquisition, an LSM510 (Zeiss, Oberkochen, Germany) laser scanning microscope using a 63x oil (1.4 NA) objective was used. To avoid overlapping signals, images were obtained by sequential excitation at 488 nm, 543 nm, and 633 nm. Images were processed using Adobe Photoshop CC 2021 software.

To examine the expression pattern of myosin heavy chain I (MyHCI) in TA muscle, an antigen retrieval step using 10mM citrate buffer at pH 6.0 was performed on transversal cryosections prior to immunostaining. Tissue sections were then permeabilized with PBS containing 0.1% Triton X-100 for 30 min, blocked in 10% normal goat serum and subsequentially incubated with the primary antibodies: rabbit monoclonal anti-slow skeletal MyHCI (1:100, Abcam, Cambridge, UK, cat. ab234431) and rat monoclonal anti-laminin-2 antibody (Lam, 1:150, Sigma-Aldrich, cat. L0663). Once washed with PBS, sections were incubated for 1 h with the secondary fluorescent antibodies: Alexa Fluor 488 AffiniPure Donkey Anti-Rabbit IgG (H+L) and Cy5 AffiniPure Donkey Anti-Rat IgG (H+L) (1:500, Jackson Immuno Research Laboratories, West Grove, PA, USA). After washing, slides were treated with the Autofluorescent Eliminator Reagent (Merck, Darmstadt, Germany), to avoid muscle fiber autofluorescence, and coverslipped with the anti-fade mounting medium Mowiol (Sigma-Aldrich). Whole TA muscle transversal section images were obtained with a laser scanning confocal microscope (Fluoview FV-1000, Olympus, Tokyo, Japan). Briefly, a sequence of single optical sections of a TA muscle mid-belly cross-section from each animal was imaged using an x60 oil immersion Olympus objective and stitched with the Fluoview FV-1000 Olympus software to create a mosaic image containing the whole TA muscle transversal section.

### Electron microscopy

For transmission electron microscopy, animals were perfused with 3% glutaraldehyde in 0.12 M phosphate buffer (0.12 M Na_2_HPO_4_ and 0.12 M NaH_2_PO_4_; pH 7.2). TA muscles were dissected and further fixated for 3 h in the same fixative. Small pieces of muscle were washed in 0.12 M phosphate buffer, fixed in 1% O_s_O_4_ for 2 h, washed again, dehydrated in increasing concentrations of acetone, and embedded in Araldite resin (Electron Microscopy Sciences, Hatfield, PA, USA). Ultrathin sections were mounted on copper grids and stained with uranyl acetate and lead citrate for examination.

For immunogold electron microscopy, animals were perfused with 3.7% PFA in 0.1 M cacodylate buffer at pH 7.4. TA muscles were dissected and further fixated for 3 h in the same fixative. Small pieces of the muscle samples were washed in 0.1 M cacodylate buffer, dehydrated at −20 °C in increasing concentrations of methanol, embedded in Lowicryl K4M (Electron Microscopy Sciences), and polymerized with ultraviolet irradiation at −20 °C for 7 days. Ultrathin sections were mounted on nickel grids and sequentially incubated with 0.1 M glycine in PBS (15 min), 5% BSA in PBS (30 min), and the primary mouse antibody anti-skeletal Myosin Heavy Chain (US Biologicals, Swampscott, MA, USA, M9850-15B) diluted (1:25) in 50 mM Tris-HCl, pH 7.6, containing 5% BSA for 2 hat 37 °C. After washing, the sections were incubated with the goat anti-mouse IgG antibody coupled to 15-nm gold particles (BioCell, Anaheim, CA, USA; diluted 1:50 in PBS containing 1% BSA) for 1 h at room temperature. Following immunogold labeling, the grids were stained with uranyl acetate.

Observations were performed with a JEOL JEM-1011 transmission electron microscope (JEOL, Tokyo, Japan) operating at 80 kV.

### Protein expression analysis

For western blotting analysis, TA muscle samples were lysed in RIPA buffer (Pierce Biotechnology, Waltham, MA, USA) supplemented with proteinase and phosphatase inhibitor cocktails (Roche, Basel, Switzerland) using a Sonic Dismembrator (Fisher Scientific), incubated on ice for 10 min, and cleared by centrifugation at 4°C for 10 min. Protein concentration from total muscle lysate supernatants was determined using BCA Protein Assay kit (Pierce Biotechnology) according to the manufactureŕs protocol. Equal amounts of protein lysate were resolved in Novex 4–20% Tris-glycine gels (Invitrogen) by SDS-PAGE electrophoresis and transferred onto Whatman Protran 0.22 µm nitrocellulose transfer membrane filters (Sigma-Aldrich) using a wet Mini Trans-Blot Cell (BIO-RAD, Hercules, CA, USA). The membranes were blocked in 5% blotting-grade milk (BIO-RAD) and probed with a mouse monoclonal antibody anti-SMN (BD Biosciences, San Jose, CA, USA; 610646) and a rabbit polyclonal antibody anti-LaminA/C (kindly donated by Prof. Larry Gerace). After extensive washing, blots were developed using specific HRP-conjugated secondary antibodies and protein bands were visualized using Western HRP Substrate (Li-COR, Lincoln, NE, U.S.A) and a C-DiGit Blot Scanner (Li-COR).

### Gene expression analysis

For qRT-PCR, TA muscles from mice, sacrificed by cervical dislocation, were quickly removed and snap frozen in liquid nitrogen. RNA was isolated with TRIzol following the manufacturer’s instructions (Invitrogen, Carlsbad) and purified with the PureLink RNA kit (Invitrogen). The concentrations of total RNA were determined using a NanoDrop ND-1000 spectrophotometer (Nanodrop Technologies, Spain). One µg of RNA was reverse-transcribed to first-strand cDNA using the High-Capacity cDNA Reverse Transcription Kit (Life Technologies) and random hexamers as primers. The expression of specific mRNAs was determined by RT-qPCR using gene-specific SYBR Green-based primers (Invitrogen). Each individual RT-qPCR assay was carried out in triplicates. The threshold cycle (Ct) for each well was determined and the results were normalized to GAPDH as a reference. Relative gene expression was calculated according to the 2^-(ΔΔCt)^ equation (Livak and Schmittgen, 2001). SYBR Green-based specific primers for murine RNAs were: for *Gapdh* 5’-AGGTCGGTGTGAACGGATTTG-3’ and 5’-TGTAGACCATGTAGTTGAGGTCA-

3’; *IL-15* 5’-GCAATGAACTGCTTTCTCCTGG-3’ and 5’-GCAGCCAGATTCTGCTACATTC-3’; *MuRF1* 5’-TACGACGTCCAGAGGGATGA-3’ and 5’-TGCCATCCGCTTGCATTAGA-3’; *Atrogin-1* 5’-CACATTCTCTCCTGGAAGGGC-3’ and 5’-TTGATAAAGTCTTGAGGGGAA-3’; *Dcn* 5’-GGGCTGGCACAGCATAAGTA-3’ and 5’-GGACAGGGTTGCCGTAAAGA-3’; *Pax7* 5’*-*GCCAAGAGGTTTATCCAGCC-3’ and 5’-AGAGGGGTGGACACTTCCAG-3’; *MyoD 5’-*TACAGTGGCGACTCAGATGC-3’ and 5’-GAGATGCGCTCCACTATGCT-3’; *Myog* 5’-AGGAGATCATTTGCTCGCGG-3’ and 5’-TTTCGTCTGGGAAGGCAACA-3’; *Mrf4* 5’-CCAACCCCAACCAGAGACTG-3’ and 5’-TTCTCTTGCTGATCCAGCCG-3’; *PGC1*□ 5’-GTCATGTGACTGGGGACTGT-3’ and 5’-GACGCCAGTCAAGCTTTTTCA-3’; *Mfn1* 5’-CCAGGTACAGATGTCACCACAG-3’ and 5’-TTGGAGAGCCGCTCATTCACCT-3’; *Mfn2 5’-*GTGGAATACGCCAGTGAGAAGC-3’ and 5’-CAACTTGCTGGCACAGATGAGC-3’; *Fndc5* 5’-GGGCAGGTGTTATAGCTCTCTT-3’ and 5’-TCATATCTTGCTGCGGAGGAG-3’.

### Morphometric analysis

Morphometric analyses of the myofiber diameter, mitochondrial area, and proportion of MyHCI-positive myofibers was performed using ImageJ software (US National Institutes of Health, Bethesda, MD, USA).

Myofiber diameter was measured on confocal images of transversal TA muscle cryosections stained with Phalloidin-FITC. The lesser myofiber diameter, defined as “the maximum diameter across the lesser aspect of the muscle fiber” described by (Dubowitz and Sewry, 2007a), was measured from at least 200 myofibers per animal.

For transversal mitochondrial area determination, transmission electron microscopy images of TA muscle ultrathin sections were used. The external perimeter of each myofiber and of its corresponding nuclei and mitochondria were delineated, and the transversal area measured. Total sarcoplasmic area occupied by mitochondria for each myofiber, excluding the nuclear area, was calculated, and relativized to percentage. At least 50 myofibers per animal were examined.

The proportion of MyHCI-positive myofibers was analyzed on entire whole TA muscle cross-section mosaic images immunostained with anti-MyHCI and anti-Lam antibodies. First, the total number of myofibers was semi-automatically determined in each muscle cross-section (one mid-belly section per TA muscle, n = 4 animals per genotype). Briefly, Lam binarized staining was used for outlining individual myofibers, and each fiber-belonging particle was analyzed. Then, the number of MyHCI-positive myofibers was manually counted and relativized to percentage.

The number of MNs was manually counted on confocal images of transversal lumbar spinal cord cryosections stained with PI. MN counts were performed on at least 10 hemispinal cord images per animal.

### Statistical analysis

Data are presented as mean ± standard deviation unless otherwise stated. For comparisons among experimental groups, two-tailed Student’s *t*-test was performed using GraphPad Prism 8.0.1 (GraphPad Software) with significance set at *p* < 0.05.

## RESULTS

### SMA myopathy in the SMN**Δ**7 mouse involves two consecutive pathological windows: preneurodegenerative and neurodegenerative stages

Several studies in mouse models of this disease have reported skeletal myofiber defects, including molecular composition alterations, mitochondrial dysfunction, and strength loss (Boyer et al., 2013a; Chemello et al., 2023; Cifuentes-Diaz et al., 2001; Mutsaers et al., 2011; Zilio et al., 2022). To explore whether SMN deficiency leads to myopathy independently of muscle denervation, we used the TA muscle, a hindlimb fast-twitch skeletal muscle that appears to be highly resistant to denervation in this disease. Indeed, in a previous study, we reported that the TA muscle of SMNΔ7 mice does not exhibit significant denervation at P8 (Cerveró et al., 2016), in contrast to what occurs in proximal muscles of the same SMA mouse model (e.g., intercostalis muscle), which shows more prominent denervation at this age (Cerveró et al., 2016; Woschitz et al., 2022).

We first examined the SMN protein expression levels in tissue lysates from the TA muscles of SMNΔ7 and WT mice by western blotting. Compared to those in age-matched WT animals, SMNΔ7 mice exhibited a strong reduction in SMN levels at all postnatal (P) ages examined (P1, P5, P10 and P14) (Fig. S1A). A similar decrease in SMN levels has been previously reported in skeletal muscles (i.e., gastrocnemius) of two different murine SMA models, *Taiwanese* and *Smn^2B/-^* mice (Groen et al., 2018). The decreased expression of SMN in muscle at early stages of the disease (i.e., P0 and P5) prompted us to examine whether SMN deficiency could provoke myopathy independent of MN loss in SMNΔ7 mice.

For this purpose, we counted the number of MN cell bodies in transverse cryosections of the spinal cord (L4 and L5 segments) stained with propidium iodide according to previously reported criteria (Mancuso et al., 2014; Riancho et al., 2015). Importantly, compared with those in WT mice, a significant decrease in the number of MNs was found in SMNΔ7 animals at P10, which progressed to reach an ∼35% reduction at P14 (Fig. S1B).

The time course of MN loss led us to establish two postnatal evolutive stages in SMNΔ7 mouse myopathy: *i)* the preneurodegenerative stage (PND, from P0 to P7), characterized by muscle SMN deficiency but with the absence of overt MN loss and myofiber denervation (Cerveró et al., 2016), and *ii)* the neurodegenerative stage (ND, from P8 to P14), characterized by both SMN deficiency and denervation-dependent (neurogenic) atrophic myofibers (Fig. S1C).

The ND stage has been the subject of extensive research on SMA myopathy in humans and mouse models of this disease (Berciano et al., 2020a; Castillo-Iglesias et al., 2019; Dubowitz and Sewry, 2007b; Fletcher et al., 2017; Jackman and Kandarian, 2004; Lunn and Wang, 2008; Prior, 2010). In the present study, we preferentially focused on the PND stage, which allows us to analyze the vulnerability of SMN-deficient myofibers before significant MN loss and muscle denervation occur. We mainly used P5 as a representative time point of the PND stage in the SMNΔ7 mouse. We examined the impact of SMN depletion on the contractile cytoskeleton of myofibrils as well as on the organelles involved in the muscle contraction-relaxation cycle and excitation-contraction coupling process, particularly the sarcoplasmic reticulum (SR), triads, and intermyofibrillar mitochondria.

### SMN deficiency during the PND stage entails nonatrophic myopathy in SMN**Δ**7 mice

To determine whether the reduced SMN levels in the PND stage were associated with motor dysfunction (Feather-Schussler and Ferguson, 2016), we performed the righting reflex test at several postnatal days (Supplementary Video 1-4). As previously reported (Berciano et al., 2020b), at P3, WT mice achieved a motor capacity of 180° rotation in less than 10 s; in contrast, SMNΔ7 mice needed more than 30 s to reach the supine position (Fig. 1A and Supplementary Video 1, 2). SMNΔ7 mice exhibited muscle weakness and hindlimb paresis during the PND stage (Supplementary Video 1, 2), which progressed to very severe paralysis at the ND stage (Supplementary Video 3, 4). These data indicate that SMN depletion is associated with motor deficits before the onset of MN death and muscle innervation loss.

**Figure 1.**
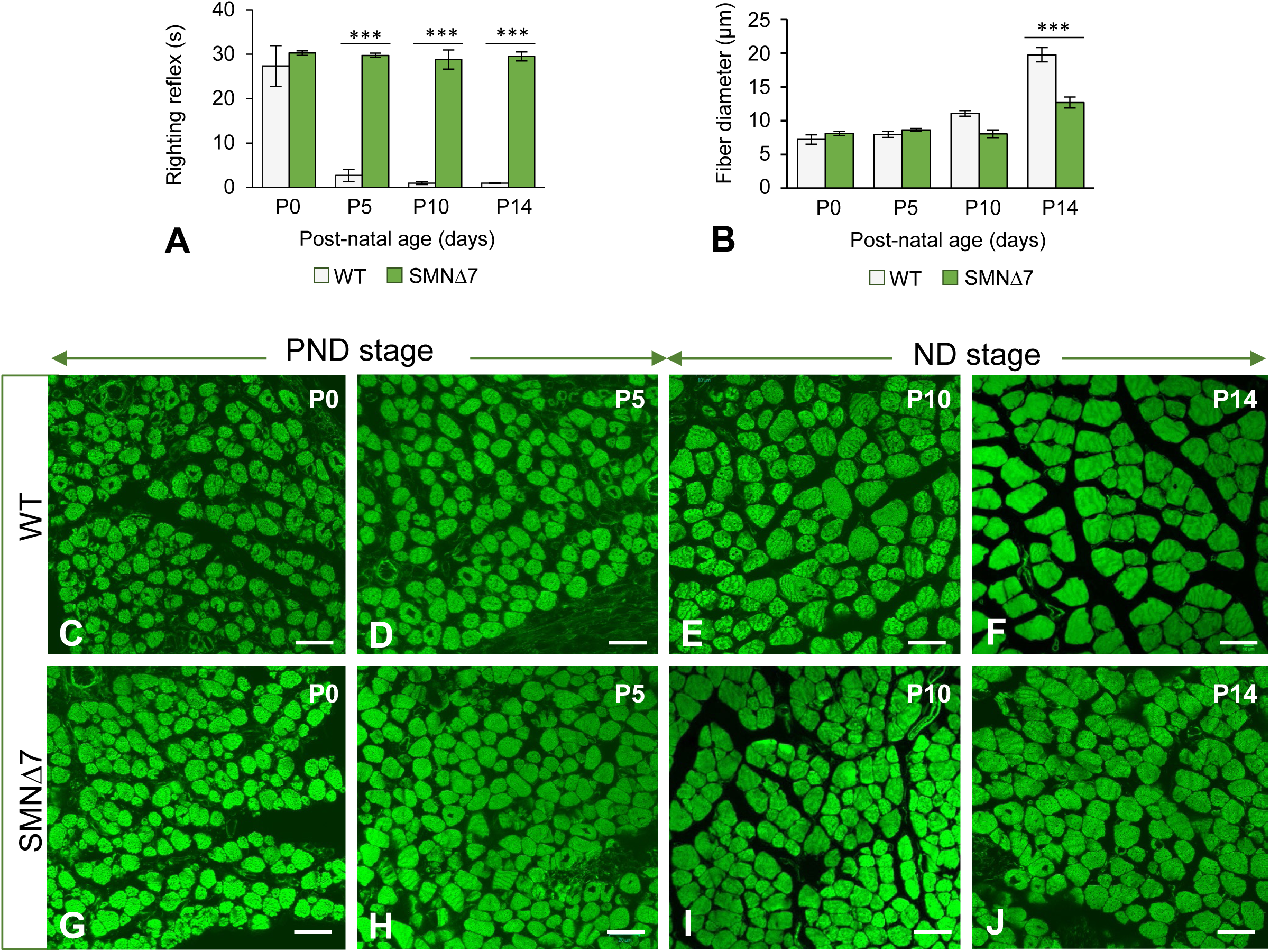
**(A)** Quantitative analysis of the righting motor reflex acquisition by WT (n=6) and SMNΔ7 (n=6) mice at the indicated postnatal ages. *p* values from WT and SMNΔ7 data comparison were 0.2516, 3.1E-10, 1.7E-10 and 5.5E-11 at P0, P5, P10 and P14, respectively. **(B)** Quantitative analysis of the mean myofiber diameters from WT (n=5) and SMNΔ7 (n=5) TA muscle at the indicated postnatal ages. At least 500 measurements per experimental group were performed using the Image J software (National Institutes of Health, USA) on transversal cryosections stained with FITC-phalloidin. *p* values from WT and SMNΔ7 data comparison were 0.0618, 0.0647, 1.5E-05 and 8.9E-06 at P0, P5, P10 and P14, respectively. ***: *p* < 0.0005. **(C-J)** Representative confocal microscopy images of transversal cryosections of TA muscle, stained with FITC-Phalloidin, used for the quantification of the mean myofiber diameter shown in panel B. Images from WT (C-F) and SMNΔ7 (G-J) mice at P0 (C, G), P5 (D, H), P10 (E, I) and P14 (F, J). Scale bar: 10µm (C-J).

Since the postnatal maturation of motor function positively correlates with the body growth rate (Chou et al., 2021), we determined the body weight of the WT and SMNΔ7 mice from P0 to P14. Compared to that of WT animals, SMNΔ7 mice exhibited a moderate but significant decrease in the growth-related gain in body weight during the PND stage (Fig. S1D). To investigate whether muscle weakness and hindlimb paresis observed at PND were accompanied by muscle atrophy, the main pathological hallmark of neurogenic SMA (Dubowitz and Sewry, 2007b; Lunn and Wang, 2008; Prior, 2010), we performed a morphometric analysis of myofiber diameter on cross sections of the TA muscle stained with FITC-phalloidin (a marker of F-actin in thin myofilaments). No significant changes in myofiber diameter were found between the TA muscles of the WT (Fig. 1B, C, D) and SMNΔ7 mice during the PND stage (Fig. 1B, G, H). Conversely, at the ND stage, a progressively significant reduction in myofiber size was observed in the SMNΔ7 mice (Fig. 1B, I, J) compared to the age-matched WT animals (Fig. 1B, E, F). These results support the view that MN loss is required for muscle atrophy to occur (Berciano et al., 2020a; Boyer et al., 2014; Cifuentes-Diaz et al., 2001; Dubowitz and Sewry, 2007b; Ling et al., 2012).

To further understand the mechanism underlying the absence of muscle atrophy in the PND stage of SMA pathology, we analyzed the expression levels of genes involved in muscle atrophy and sarcopenia in the TA muscle at P5 by RTLqPCR. First, we explored the expression of *IL-15*, a myokine that stimulates myofilament protein biosynthesis and is usually suppressed in individuals with muscular atrophy (O’Leary et al., 2017). No significant changes in the *IL-15* levels were observed in muscles from SMNΔ7 mice relative to those from WT animals (Fig. 2A). We subsequently analyzed two atrogenes, *MuRF1* and *MAFbx/atrogin-1 (atrogin-1)*, which are biomarkers of muscle atrophy (Bodine and Baehr, 2014). Both genes encode muscle-specific E3 ubiquitin ligases that are upregulated in different murine SMA models under conditions of atrophy induction (Altun et al., 2010; Bodine and Baehr, 2014; Deguise et al., 2016; Jackman and Kandarian, 2004). Importantly, compared with those in the WT samples, no significant changes in the expression levels of either of the two atrogenes were observed in the TA muscle from SMNΔ7 mice (Fig. 2B-C). Later, we investigated the expression of *Dcn*, which encodes decorin, a small leucine-rich proteoglycan that prevents muscle atrophy by inhibiting the action of myostatin (El Shafey et al., 2016) and negatively regulating *atrogin1* and *MuRF1* expression (Kanzleiter et al., 2014; Lee and Jun, 2019; Marshall et al., 2008). Interestingly, our results revealed a marked increase (10-fold increase) in *Dcn* expression in SMNΔ7 mice compared to that in age-matched WT animals (Fig. 2D). This finding is consistent with a reactive *Dcn* upregulation response to SMN depletion to prevent muscle atrophy in SMNΔ7 mice during the PND stage.

**Figure 2.**
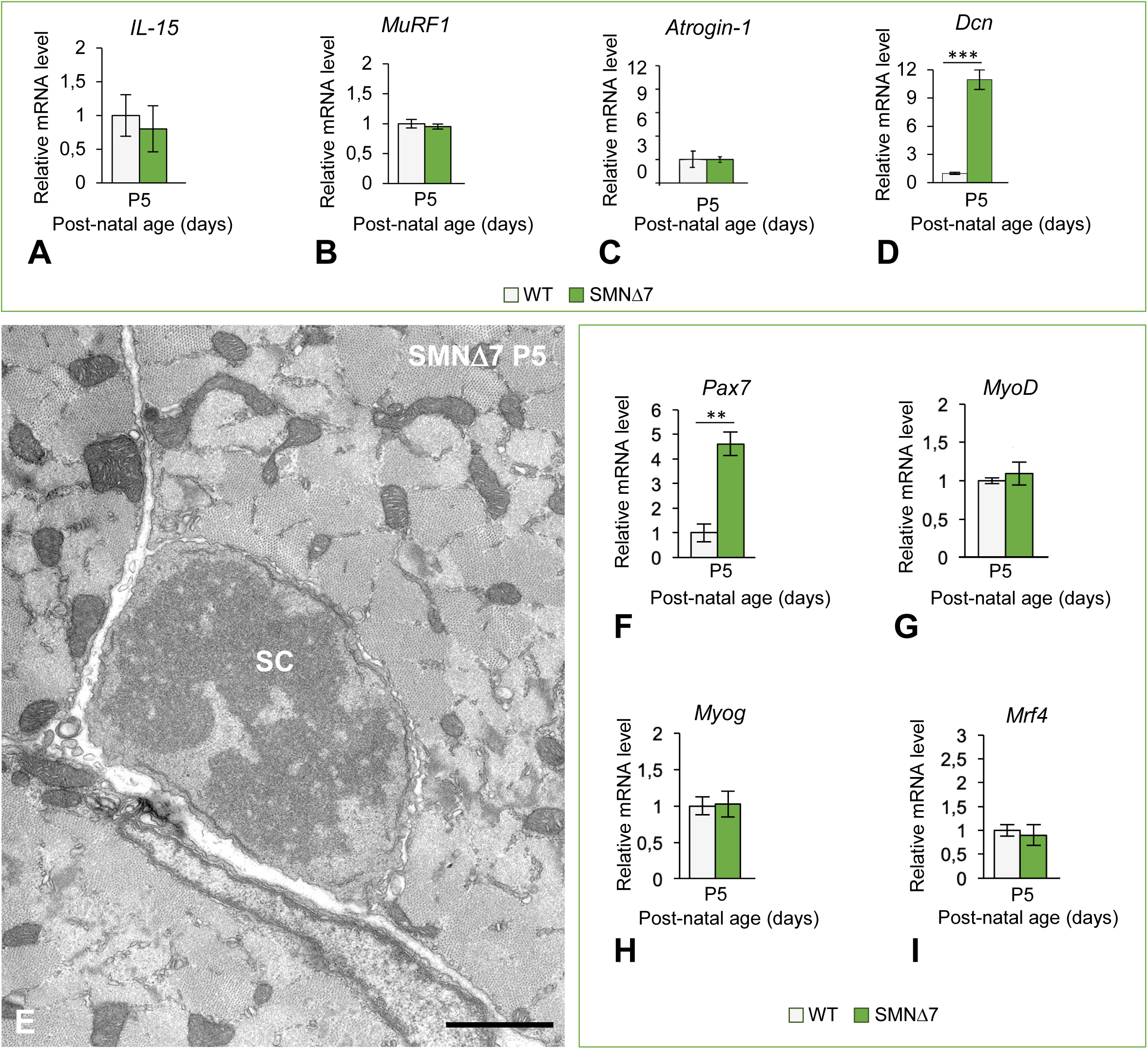
**(A-D)** qRT-PCR determination of the normalized expression levels of *IL-15* (A), *MuRF1* (B), *Atrogin-1* (C) and *Dcn* (D) mRNAs in TA extracts from P5 SMNΔ7 mice (n=5) relativized to age-matched WT animals (n=3). *p* values from WT and SMNΔ7 data comparison were: 0.5588 for *IL-15*, 0.4011 for *MuRF1*, 0.4919 for *Atrogin-1* and 0.0005 for *Dcn*. ***: *p* < 0.0005. **(E)** Electron micrograph illustrating a muscle satellite cell (SC) in telophase closely attached to a myofiber in transversal section. Scale bar: 2mm. **(F-I)** qRT-PCR determination of normalized mRNA expression levels of *Pax7* (F), *MyoD* (G), *Myog* (H) and *Mrf4* (I) in RNA extracts of TA muscle samples from SMNΔ7 mice (n=5) and age-matched WT animals (n=3) at P5. Note in SMNΔ7 samples the approximately five-fold increase of *Pax7* mRNA expression at the PND stage. *p* values from WT and SMNΔ7 data comparison were: 0.00296 for *Pax7,* 0.3924 for *MyoD,* 0.7655 for *Myog* and 0.5049 for *Mrf4*. **: *p* < 0.005.

Since decorin also regulates muscle mass by modulating postnatal myogenesis (Kishioka et al., 2008; Li et al., 2007), we analyzed whether the postnatal expression of some myogenic regulatory factors was preserved in the TA muscle of SMNΔ7 mice at the PND stage. The expression of Pax7, a key transcription factor for satellite cells involved in regenerative myogenesis (Collins et al., 2009), was significantly greater in SMNΔ7 muscles at P5 than in age-matched WT tissue samples (Fig. 2F). Consistent with the overexpression of *Pax7,* we detected mitosis in satellite cells at the PND stage (Fig. 2E). Moreover, no changes in the expression levels of three key myogenic regulatory factors, *MyoD1, Myog* (myogenin) or *Mrf4,* which are required for skeletal muscle specification, differentiation, and maturation (Kassar-Duchossoy et al., 2004; Valdez et al., 2000), were observed when SMNΔ7 and WT TA muscle samples were compared at P5 (Fig. 2 G-I). We believe that the preserved expression of these myogenic regulatory factors during the PND stage could contribute to the normal growth of myofibers and, consequently, to the absence of muscle atrophy.

### SMN deficiency induces focal and segmental lesions in myofibrils with abnormal accumulation of F-actin and sarcomere disruption in the absence of MN loss

In a previous study of human type I SMA (Berciano et al., 2020a), we demonstrated the existence of focal and segmental lesions in nonatrophic (innervated) myofibers with different degrees of sarcomere disruption, suggesting an intrinsic effect of SMN depletion on sarcomere architecture. This prompted us to address whether such myofiber lesions also occur during the PND stage in SMNΔ7 mice. For this purpose, we performed confocal and electron microscopy analyses.

For confocal microscopy, we used longitudinal cryosections of the TA muscle of WT and SMNΔ7 mice at the PND stage (P0 and P5) stained with phalloidin-FITC. At both ages examined, the WT myofibers exhibited the typical striated morphology with properly aligned sarcomeric I- and A-bands (Fig. 3A). In contrast, numerous SMNΔ7 myofibers exhibited focal lesions during this stage (Fig. 3B-D). The most striking alteration was the presence of unstructured myofiber areas with a loss of cross-striation and abnormal accumulation of fluorescent F-actin signals, which were already detected at P0 (Fig. 3B-D). Another common finding was the presence of myofiber segments with overcontracted “minisarcomeres”. Both types of lesions progressed from the PND stage to the ND stage (Fig. 3E). Interestingly, myofiber lesions were commonly flanked by well-preserved myofibrils with a sharp transition between normal and damaged sarcoplasmic areas. Moreover, double fluorescent labeling with phalloidin-FITC and a nuclear marker of TA cryosections at P5 revealed the typical peripheral positioning of myonuclei in WT myofibers (Fig. 3F), as well as the presence of some central myonuclei and the frequent spatial association of areas of sarcomere disruption with myonuclei in SMNΔ7 myofibers (Fig. 3G, H).

**Figure 3.**
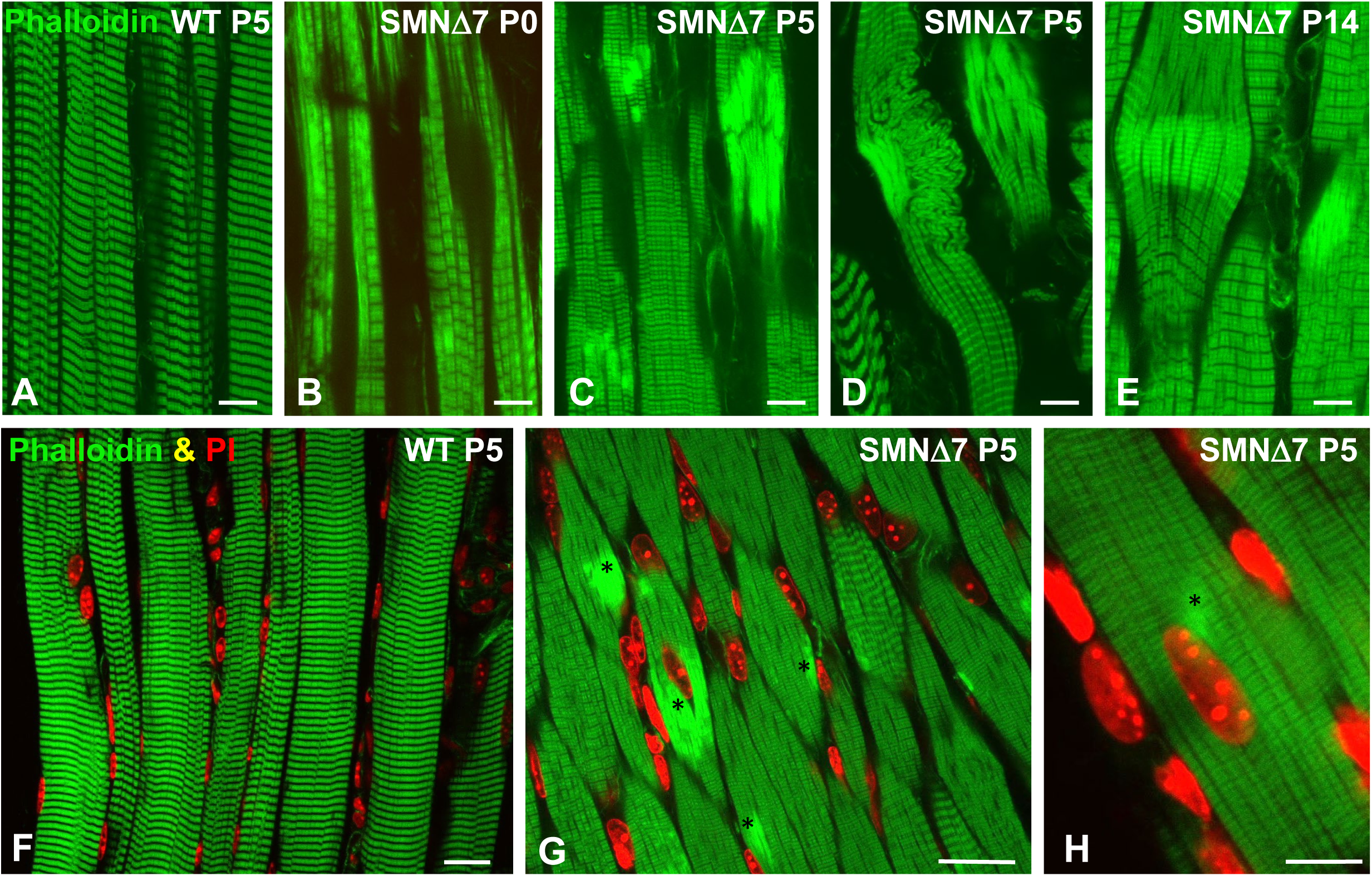
**(A-E)** Representative confocal images of longitudinal cryosections of the TA muscle stained with FITC-phalloidin from WT at P5 (A) and SMNΔ7 mice at P0 (B), P5 (C-D) and P14 (E). Note in SMNΔ7 mice images the presence of myofiber regions with disruption of cross striation and aberrant accumulations of F-actin showing bright Phalloidin-FITC fluorescent signal. **(F-H)** Double staining to label actin filaments (FITC-phalloidin, green channel) and nuclei (DAPI, red channel) in WT (F) and SMNΔ7 (G-H) myofibers at P5. Note the typical cross striation and peripheral positioning of myonuclei in WT myofibers (F), and the presence of several bright foci of F-actin accumulations, some of them closely associated with central or peripheral myonuclei (asterisks) in SMNΔ7 myofibers (G, H). Scale bars: 20mm (A-G) and 10mm (H).

Electron microscopy analysis of longitudinal myofiber sections at the PND stage confirmed the presence of SMNΔ7 myofibers with focal and segmental lesions that coexisted with normal unaltered myofibers. At P5, whereas WT myofibers exhibited regularly spaced and aligned myofibrils with sarcomeres clearly delimited by well-defined Z-discs (Fig. 4A), some SMNΔ7 myofibers displayed overt structural alterations (Fig. 4B-F). These included the misalignment of myofibrils (Fig. 4B); small focal areas of sarcomere disruption, which involved one or a very few sarcomeres usually located adjacent to myonuclei (Fig. 4C, D); and larger segmental areas of sarcomere disruption, which affected extensive myofiber segments with sarcoplasmic integrity loss commonly found during the ND stage (Fig. 4E-H). Ultrastructural analysis also confirmed the sharp transition between affected and preserved sarcomeres (Fig. 4E, F).

**Figure 4.**
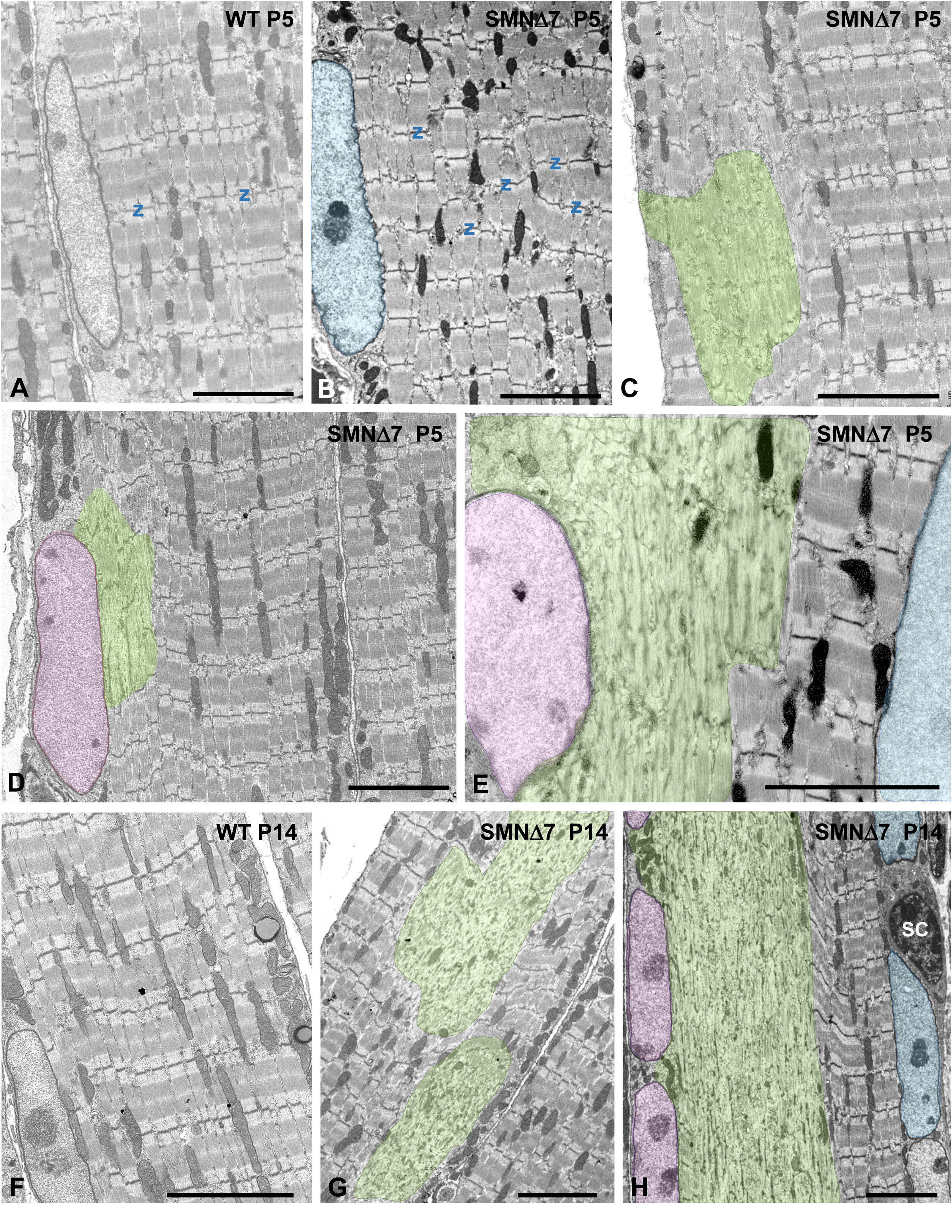
**(A-H)** Representative electron micrographs of longitudinal sections of TA myofibers from WT (A, F) and SMNΔ7 (B-E, G, H) mice. **(A**, **F)** Normal fine structure and organization of sarcomeres in WT myofibers at P5 (A) and P14 (F). **(B-E, G-H)** Ultrastructural alterations of the contractile machinery with misalignment of Z discs (Z) and presence of local sarcoplasmic areas of variable size with disruption of sarcomere architecture (transparented in green) at P5 (B-E) and P14 (G, H). Note the close spatial association of some myonuclei (transparented in pink) with areas of myofiber lesions. **(E, H)** Potential influence of myonuclear domains in the spatial distribution of myofiber lesions: in the same myofiber segment coexist peripheral myonuclei (transparented in pink) associated with areas of sarcomere disruption whereas, in the opposite side, other myonuclei (transparented in blue) associated with properly structured sarcomeres. Scale bars: 5mm (A-H).

The frequent spatial association we observed between sarcoplasmic areas of myofiber lesions and myonuclei (Fig. 4D) suggested dysfunction of specific “myonuclear domains”, which are considered key determinants of regulation and function in multinucleated skeletal myofibers (Bagley et al., 2023; Gundersen and Bruusgaard, 2008; van der Meer et al., 2011). Accordingly, in longitudinal sections of SMNΔ7 myofibers, we observed a sarcoplasmic band indicating sarcomere disruption adjacent to a row of peripheral myonuclei, whereas on the opposite side of the same myofiber, other myonuclei associated with perfectly preserved sarcomeres could be observed (Fig. 4E, H).

High-magnification electron microscopy showed that focal lesions in the SMNΔ7 muscles were characterized by the loss of the regular pattern of A- and I-bands, which appeared blurred with distorted Z-discs (Fig. 5A, B) and were accompanied by the disassembly of thick myosin myofilaments (Fig. 5C). At larger segmental areas of sarcomere disruption, the abnormal accumulation of disordered thin actin myofilaments, reduction of thick filaments and absence of Z-discs were observed (Fig. 5D). Immunogold electron microscopy for myosin revealed the regular structure of the immunolabeled A-band of thick myofilaments with well-aligned sarcomeres in WT mice at P5 (Fig. 5E). In contrast, several alterations, including misaligned myofibrils (Fig. 5F) and different degrees of sarcomeric disarray with loss of immunogold-labeled thick myofilaments, were observed in SMNΔ7 myofibers at the PND stage (Fig. 5G, H). This ultrastructural analysis also confirmed the presence of some myofiber segments exhibiting repeats of “minisarcomeres” at the PND stage, with striking and strictly regular shortening flanked by well-organized myofibrils (Fig. 6A, C). Unlike in WT myofibers, in which the intermyofibrillar spaces were preserved (Fig. 6B), SMNΔ7 myofibers contained areas of overcontracted minisarcomeres lacking the I-band and lacking well-defined boundaries between the myofibrils (Fig. 6C).

**Figure 5.**
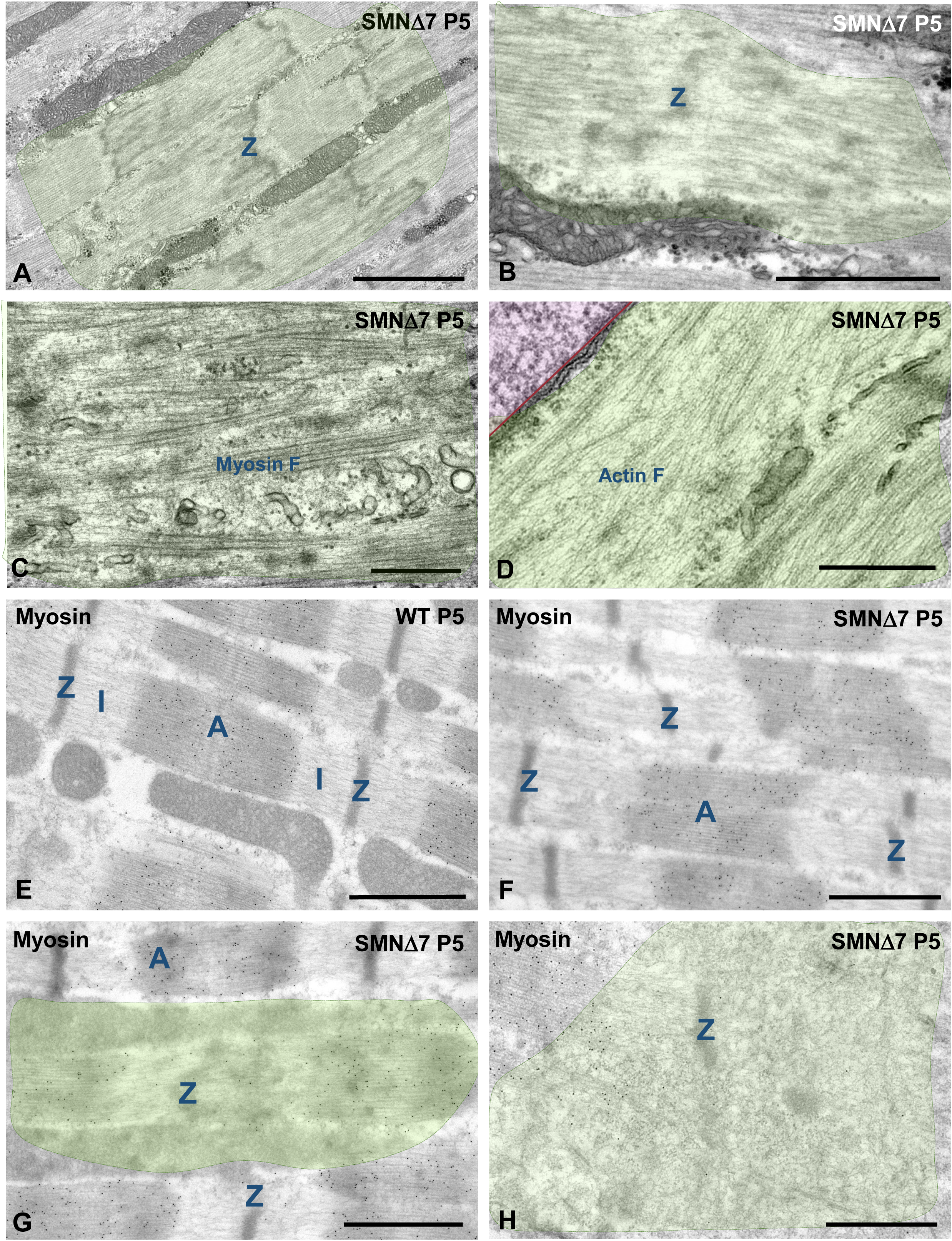
**(A-D)** Detail at high magnification of cytoskeletal alterations in SMNΔ7 myofibers at P5. **(A, B)** Disassembly of the sarcomere architecture with loss of the banding pattern and disruption of Z-discs. **(C)** Disarray of myosin thick myofilaments. **(D)** Aberrant accumulations of actin thin myofilaments. **(E-H)** Immunogold electron microscopy analysis for the detection of myosin in myofibrils from WT (E) and SMNΔ7 (F-H) mice at P5. **(E)** Gold particles of myosin immunoreactivity specifically decorate thick myofilaments in the properly aligned A-bands of WT sarcomeres. **(F-H)** In SMNΔ7 myofibers there were misalignment of A-band (F) and disarray and loss of myosin-labeled thick myofilaments in sarcoplasmic areas of unstructured sarcomeres (G-H). Scale bars: 2mm (A); 1mm (B and F-H) and 500nm (C-D).

**Figure 6.**
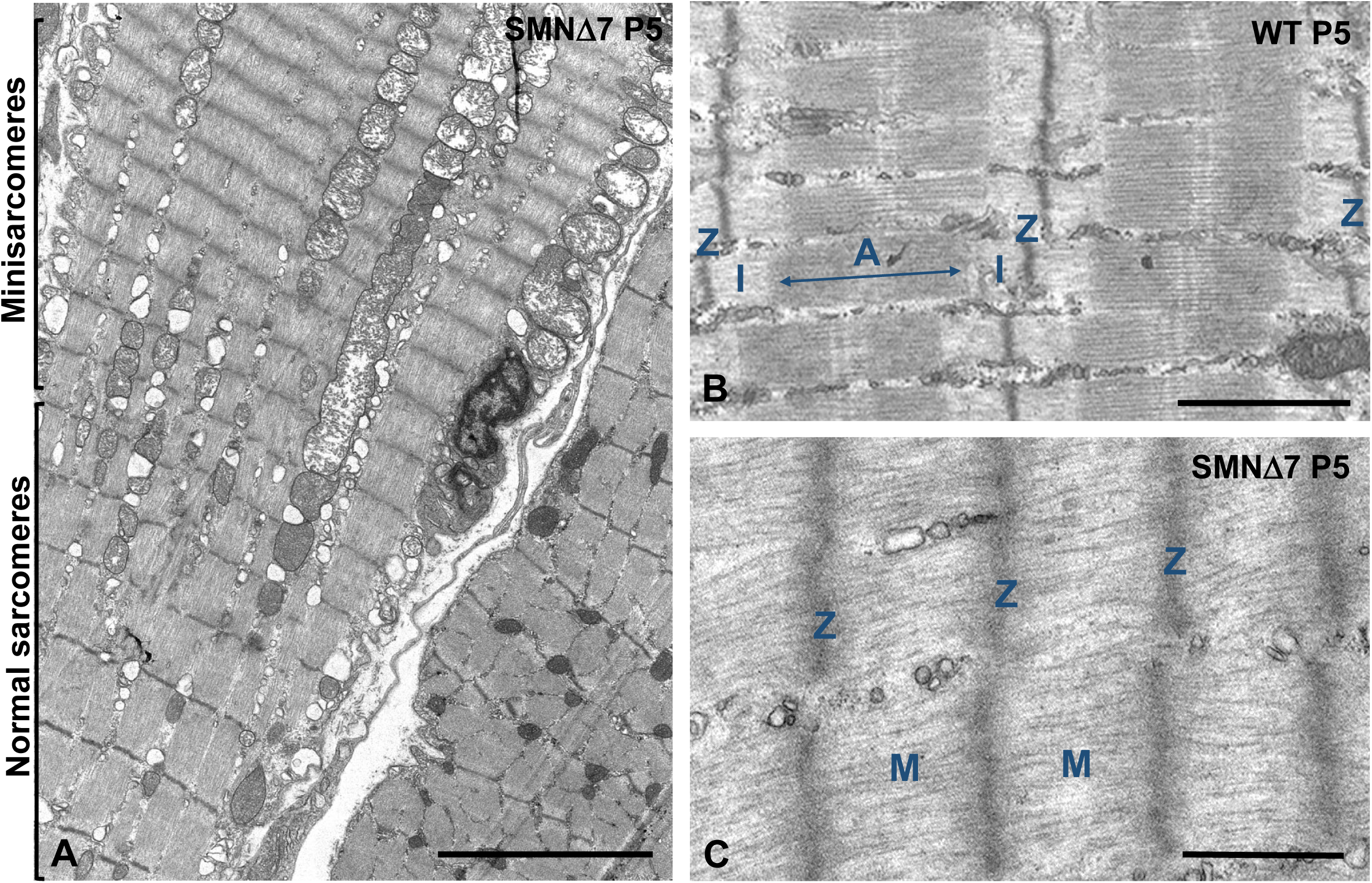
Electron micrographs of minisarcomeres in a SMNΔ7 myofiber at P5. **(A)** Panoramic view of the longitudinal section of a myofiber illustrating a segment with shortened minisarcomeres juxtaposed to another segment with normal sarcomeres. Note the sharp transition between these two myofiber segments and the swelling of some mitochondria. **(B-C)** Detail of the comparative ultrastructure between normal sarcomeres from a WT myofiber (B), with the A-band flanked by two hemi-I-bands, and shortened minisarcomeres from a SMNΔ7 myofiber (C), with absence of I-bands and thick myofilaments that are anchored directly into Z-disc. Z: Z-disc; A: A-band; I: hemi-I band. Scale bars: 5mm (A) and 1mm (B-C).

In conclusion, in SMN-deficient muscles, disturbance of the myofibrillar contractile apparatus, abnormal accumulation of actin thin myofilaments and sarcomere disruption, together with the formation of minisarcomeres, could contribute to motor impairment in SMNΔ7 mice during the PND stage.

### SMN deficiency induces sarcoplasmic reticulum (SR) and triad disruption during the PND stage

The SR, a key regulator of cytosolic Ca^2+^ homeostasis in myofibers, forms an intricate network of elongated tubules that are arranged around each sarcomere and end in dilated terminal cisterns (for a review, see (Rossi et al., 2022). The junction of two terminal cisterns with a transverse tubule (T-tubule) of the sarcolemma composes the “triad”, the structure responsible for excitation-contraction coupling (Ogata and Yamasaki, 1997; Rossi et al., 2022). Considering the key role of these mechanisms in myofiber function, we wondered whether alterations in the SR or triad could contribute to SMNΔ7 myopathy in mice at the PND stage.

Ultrastructural analysis of longitudinal sections of WT myofibers at the PND stage revealed the regular organization of axial SR tubules along the sarcomeres, as did the typical triads located at the junction between the A- and I-bands. (Fig. 7A). Moreover, intermyofibrillar mitochondria were commonly observed in association with the SR, particularly at the level of the I-band, near the Z-disc (Fig. S2A). In contrast, prominent alterations in the SR and triad density were found in TA myofibers from SMNΔ7 mice at the PND stage. In longitudinal myofiber sections, sarcoplasmic areas with misaligned myofibrils frequently showed dysmorphic SR features. These included dilations of the SR, especially of the terminal cisterns of triads (Fig. 7B, C), and, occasionally, SR swelling that eventually led to vacuolar degeneration of the sarcoplasm (Fig. 7D, E). However, the morphology of intermyofibrillar mitochondria attached to the abnormally dilated SR cistern was unaffected (Fig. 7B-D). Additionally, in transverse sections of myofibers, swelled SR cisterns were not found delineating SMNΔ7 myofibrils (Fig. 7G), in contrast with the normal appearance of SR-tubule networks, which wrap WT myofibrils (Fig. 7F).

**Figure 7.**
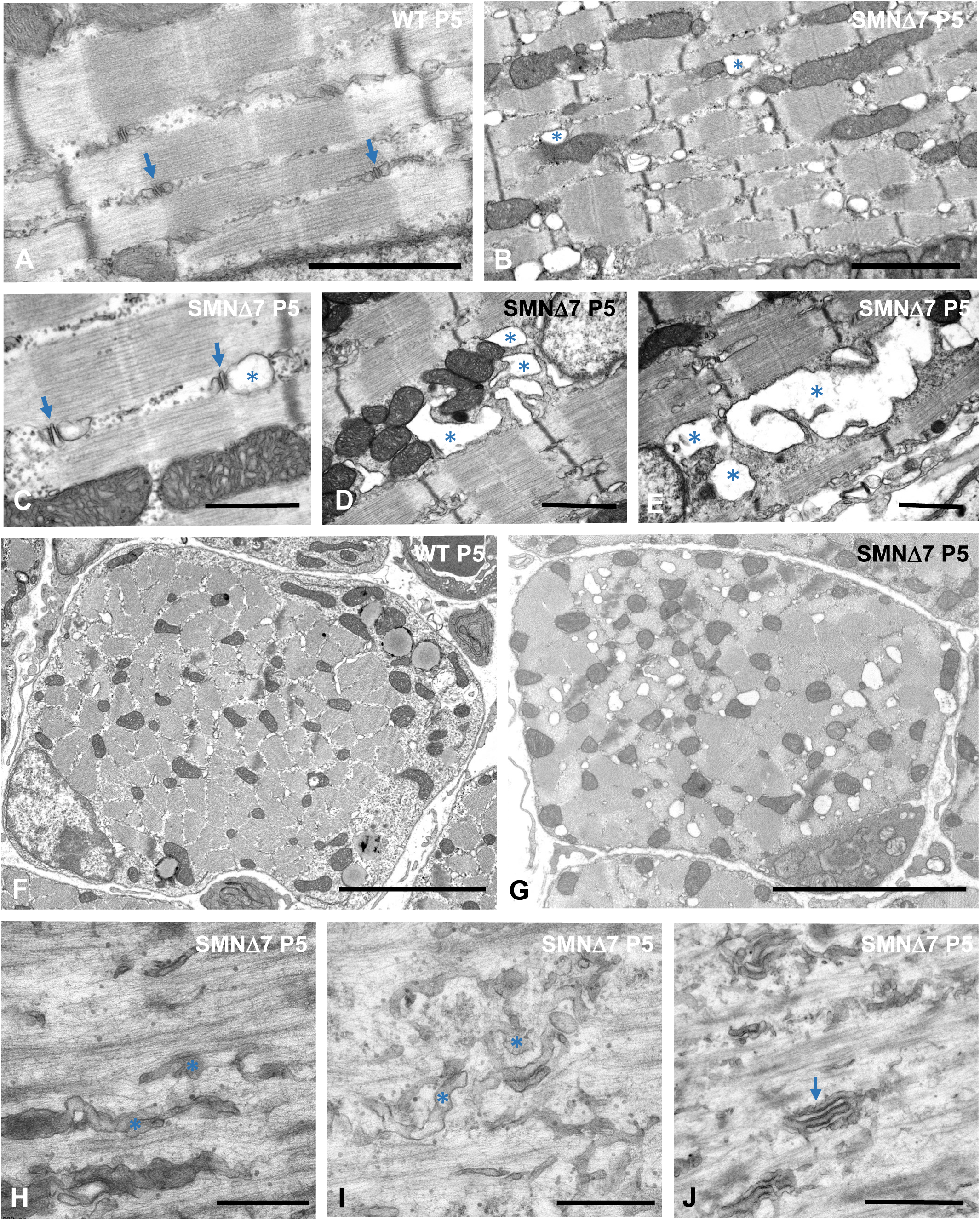
**(A-J)** Ultrastructural organization of SR and triads in WT (A, F) and SMNΔ7 (B-J; G-J) mice myofibers at P5. **(A)** Typical structure of triads, composed of a T-tubule flanked by two terminal SR cisterns (arrows), and intermyofibrillar longitudinal tubules of the SR network. **(B, C)** Dilation of SR cisterns associated with mitochondria (asterisks) and with the triads (arrows). Note the well-preserved fine structure of mitochondria. **(D, E)** Representative electron microscopy images of vacuolar degeneration affecting intermyofibrillar SR cisterns (asterisks). Note in panel D that some vacuoles directly interact with mitochondria. **(F, G)** Transversal sections of WT (F) and SMNΔ7 (G) myofibers illustrating cross-sectioned myofibrils surrounded by a network of SR tubules in the WT sample and the presence of numerous dilated cisterns of SR in the SMNΔ7 myofiber. **(H-J)** High magnification electron micrographs of sarcoplasmic areas of sarcomere disruption illustrating the disarray of SR tubules (asterisks) and triads (arrow). Scale bars: 1mm (A-E and J), 5mm (F-G) and 500nm (H, I).

Importantly, in SMNΔ7 myofiber areas with sarcomere disruption and aberrant accumulation of actin myofilaments, the SR displayed tortuous tubules with disarrangement or fragmentation of the tubular network. Furthermore, the triads appeared dysmorphic, disassembled, or unrecognized (Fig. 7H-J) and were difficult to visualize in regions with minisarcomeres (Fig. 6C).

In conclusion, the severe structural alterations in the SR and triads observed in sarcomere disrupted areas of SMNΔ7 myofibers could cause dysfunction of excitation-contraction coupling mechanisms and an imbalance in Ca^2+^ homeostasis. This could impede correct nerve impulse transmission, contributing to the primary myopathy found in the PND stage of the SMA.

### SMN deficiency induces changes in myofiber mitochondrial organization and dynamics

Previous studies have reported impaired muscle mitochondrial function in samples from patients and animal models of SMA (Berger et al., 2003; James et al., 2018; Ripolone et al., 2015; Zilio et al., 2022). Moreover, SMN depletion has been shown to cause mitochondrial dysfunction in the C2C12 cell line and in human induced pluripotent stem cells, which is dependent on the downregulation of the microRNAs (miRs) miR-1 and miR-206 (Ikenaka et al., 2023). In addition, the accumulation of dysfunctional mitochondria has been demonstrated in myofibers from a muscle-inducible *Smn1* knockout mouse model (Chemello et al., 2023). These data prompted us to study whether SMN deficiency in SMNΔ7 mice has an impact on the mitochondrial organization of TA myofibers at the PND stage (P5). Skeletal myofibers contain two distinct populations of mitochondria: subsarcolemmal mitochondria, which supply ATP for gene transcription and membrane transport, and intermyofibrillar mitochondria, which provide ATP for muscle contraction (Figs. 4A, S3A). Both mitochondrial populations associate structurally and functionally with the SR and participate in Ca^2+^ homeostasis (for review see (Dong and Tsai, 2023; Willingham et al., 2021).

MitoTracker staining of transverse cryosections of both WT and SMNΔ7 TA muscle samples revealed a predominant pattern of mitochondria-rich (oxidative) myofibers at P5 (Fig. 8A, B). The MitoTracker fluorescent signal appeared as intermyofribillar foci, which corresponded to individual mitochondria, and subsarcolemmic aggregates (Fig. 8A, B). Determination of the overall MitoTracker fluorescence signal by densitometric analysis of each myofiber showed significant differences between the WT and SMNΔ7 TA samples (Fig. 8C), suggesting that SMNΔ7 myofibers contain more mitochondria than the WT myofiber.

**Figure 8.**
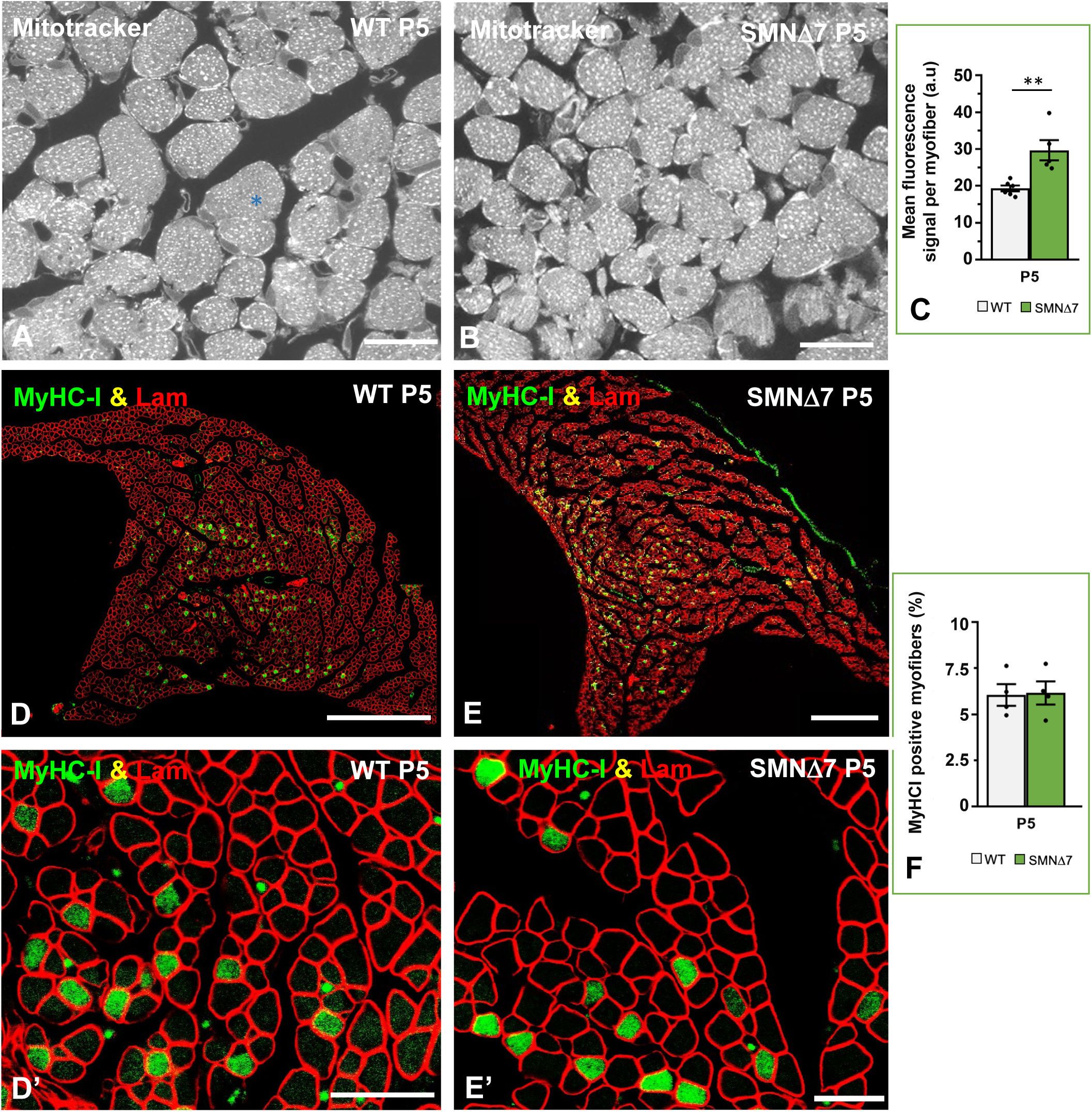
**(A-B)** Cross-cryosections of TA muscle stained with Mitotracker to analyze the mitochondrial content in WT and SMNΔ7 myofibers at P5. The asterisk indicates a fast glycolytic type II myofibers. **(C)** Determination of the mean fluorescence intensity of MitoTracker signal per myofiber, using confocal images obtained from cross-cryosections of TA muscle from WT (n=3) and SMNΔ7 (n=3) at P5 stained with Mitotracker. *p* value from WT and SMNΔ7 data comparison was 0.0034. **(D-F)** Representative confocal mosaic images of TA muscle mid-belly transversal sections from WT and SMNΔ7 mice at P5, double immunostained using antibodies against myosin heavy chain I (MyHC-I, green), for slow-type I myofibers, and laminin (Lam, red), for myofiber contour visualization. Note the non-uniform distribution of MyHC-I positive myofibers along the TA muscle. **(D’-E’)** Higher magnification views of immunolabeled TA myofibers of the different experimental conditions. **(F)** Mean percentage of MyHC-I positive fibers in TA muscle from WT and SMNΔ7 mice at P5. All myofibers of a mid-belly transversal section of TA muscle from 4 animals per experimental condition were analyzed. *p* value from WT and SMNΔ7 data comparison was 0.9023. In all graphs (C and F), values are shown as mean±SEM, and unpaired Student’s t test (C) and two-way analysis of variance (Bonferroni’s *post hoc* test) (F) was used for statistical analysis; **: *p* < 0.005. Scale bar: 10μm (A, B) 250 μm (D, E) and 25 μm (D’, E’).

Next, we investigated whether low SMN levels influence the expression of low skeletal myosin heavy chain I (MyHC-I). The postnatal adaptation of muscles to specialized contractile properties requires the expression of different isoforms of the myosin heavy chain. Thus, the embryonic and neonatal MyHC isoforms are progressively replaced by adult isoforms I (slow type), IIA, IIX/IID, and IIB (fast type) to meet the specific functional requirement of each muscle (Schiaffino, 2018; Schiaffino and Reggiani, 1996). To explore whether the greater number of mitochondria found in SMNΔ7 TA muscles was associated with a change in the myosin profile with a subsequent increase in slow-twitch, oxidative myofiber expression, we performed an immunocytochemical analysis of slow-type MyHC. On cross sections of TA muscle coimmunostained for MyHC-I and laminin (basal lamina marker), we found no significant changes in the percentage of MyHC-I-positive myofibers between WT and SMNΔ7 samples at the PND stage (Fig. 8D-F). This indicates that reduced levels of SMN in SMNΔ7 mouse myofibers induce changes in the mitochondrial compartment that are independent of the expression levels of the slow skeletal MyHC-I protein.

Ultrastructural morphometric analysis of transverse sections of TA myofibers at the PND stage (P5) confirmed that both the mitochondrial area and the proportion of the sarcoplasmic area occupied by mitochondria were significantly greater in SMNΔ7 muscle than in WT tissue (Fig. 9A-C). To confirm whether the enlargement of the mitochondrial compartment in SMNΔ7 myofibers correlated with changes in the expression of the master regulator of mitochondrial biogenesis *PGC-1a* (proliferator-activated receptor-g coactivator-1-a) (Chen et al., 2023), we performed qRTLPCR analysis. The results revealed a significant increase in *PGC-1a* transcript levels in the SMNΔ7 TA muscle at the PND stage relative to those in the WT samples (Fig. 9E). It is well established that PGC-1a activates the downstream factor FDCN5 (fibronectin type III domain containing 5), which is cleaved in the sarcolemma of skeletal myofibers to give rise to the myokine irisin (Boström et al., 2012; Chen et al., 2017; He et al., 2020; Reza et al., 2017). Due to the close relationship of the PGC-1a/FDCN5/irisin signaling pathway with genes and proteins that regulate mitochondrial biogenesis (Chen et al., 2017; He et al., 2020; Srinivasa et al., 2016), we analyzed whether the overexpression of *PGC-1a* was associated with changes in *Fdcn5* gene expression. By qRTLPCR, we found that the expression of *Fdcn5* transcripts was greater in SMNΔ7 TA muscles than in WT samples (Fig. 9F). Collectively, these findings suggest that SMN deficiency in myofibers promotes the overactivation of mitochondrial biogenesis during the PND stage.

**Figure 9.**
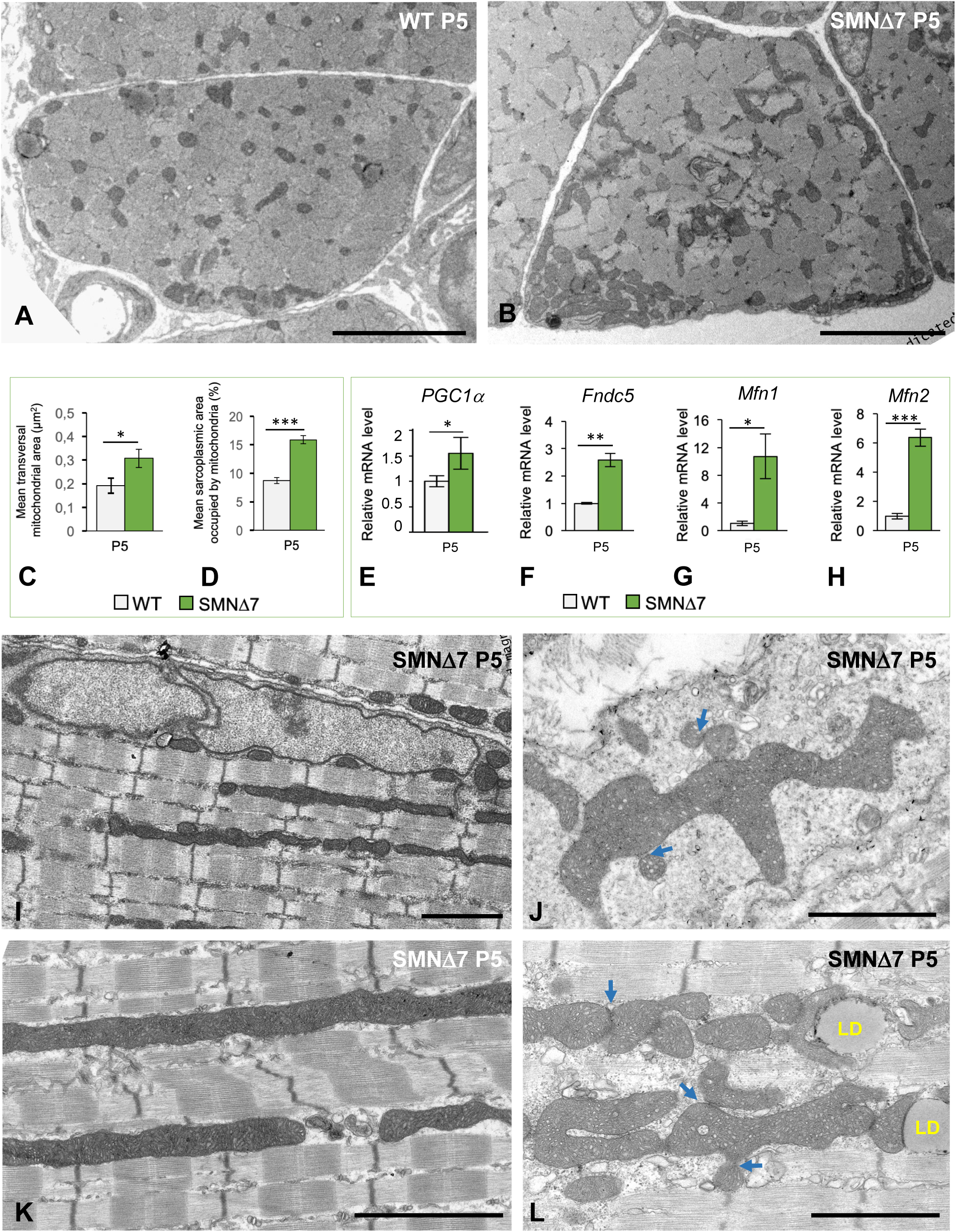
**(A, B)** Electron micrographs of cross-sectioned myofibers of TA muscle from WT (A) and SMNΔ7 (B) mice at P5. Note the accumulation of subsarcolemmal mitochondria in the SMNΔ7 myofiber. **(C, D)** Quantitative analysis on electron micrographs from transversal TA myofiber sections from WT and SMNΔ7 mice at P5. The mean mitochondrial area (C) and the sarcoplasmic area occupied by mitochondria (D) were measured from at least 50 myofibers of each genotype using the Image J software. *p* values from WT and SMNΔ7 data comparison were: 0.016 for the mean transversal mitochondrial area and 0.00015 for the mean sarcoplasmic area occupied by mitochondria. **(E-H)** qRT-PCR analysis of the expression levels of *PCG1*α*, Fndc5, Mfn1 and Mfn2* in TA muscle samples from WT (n=3) and SMNΔ7 mice (n=5) at P5. Note the significant increase of these gene transcripts in SMNΔ7 RNA extracts relative to age matched WT samples. *p* values from WT and SMNΔ7 data comparison were: 0.0041 for *PCG1*α, 0.0316 for *Fndc5,* 0.0428 for *Mfn1*, and 0.0065 for *Mfn2.* **(I, L)** Ultrastructural changes in the mitochondrial phenotype of TA myofibers from SMNΔ7 mice at P5. Note that the fine structure of mitochondria in WT myofibers at P5 is shown in Supplementary Fig. 3A. **(I, K)** Presence of very long rod-like intermyofibrillar mitochondria in SMA myofibers. **(J, L)** Clusters of subsarcolemmal (J) and intermyofibrillar (L) mitochondria illustrating the interactome (organelle contact) between mitochondria as well as between mitochondria and lipid droplets (LD). Arrows indicate potential sites of mitochondrial fusion. Scale bar: 5 μm (A, B) and 2 μm (I, J, K, L).

To further understand whether the increase in the mitochondrial compartment in SMNΔ7 myofibers correlated with the increase in mitochondrial fusion dynamics, we performed qRTLPCR on the *Mfn1* and *Mfn2* genes. These genes encode mitofusin proteins 1 and 2, respectively, two mitochondrial outer membrane GTPases that mediate mitochondrial clustering and fusion (Romanello and Sandri, 2021). Our qRTLPCR results demonstrated that both *Mfn1* and *Mfn2* were significantly upregulated in the SMNΔ7 TA muscle at the PND stage (Fig. 9G, H). Consistent with the increase in the expression of mitofusin genes, ultrastructural analysis of SMNΔ7 myofibers at P5 revealed the presence of abnormally long rod-like intermyofribillar mitochondria extending to several sarcomeres (Fig. 9I, K); these coexisted with shorter mitochondria commonly found in WT myofibers (Figs. 4A, S3A). Moreover, clusters of irregularly shaped and interacting mitochondria exhibiting outer membranes in close contact were observed at both the intermyofibrillar and subsarcolemmal regions. This finding suggested that a fusion process occurs in mitochondria in the SMA during the PND stage. (Figs. 9J, L; S3B). Notably, there was also spatial interaction between mitochondria and dilated SR cisterns (Fig. S2B, C). Myofibers harboring mitochondrial alterations, particularly swelling of the matrix with distorted cristae, were occasionally found at this stage (Fig. 6A).

A previous study showed that an increase in mitochondrial size by fusion prevents the elimination of mitochondria via mitophagy (Ojaimi et al., 2022). Consistent with this view, mitophagy and mitolysosomes were not detected via ultrastructural analysis. A paucity of lysosomes in SMNΔ7 mouse myofibers at the PND stage was also observed (Fig. S2B and D). Collectively, these results suggest that SMN deficiency induces dysfunction of the mitochondrial dynamics of fusion/fission in myofibers from SMNΔ7 mice.

To study the sarcoplasmic localization of mitochondria in SMNΔ7 mouse myofibers, double fluorescent cytochemical staining with MitoTracker and Phalloidin-FITC was performed. Severe loss of intermyofibrillar mitochondria in areas with abnormal accumulation of F-actin and sarcomere loss was observed at the PND stage (Fig. 10A-C). Electron microscopy analysis confirmed that mitochondria were absent or rarely found in areas of sarcomere disruption (Fig. 10D, E). The depletion of intermyofibrillar mitochondria in areas of sarcomere loss is consistent with the reduced ATP demand for myofiber contraction.

**Figure 10.**
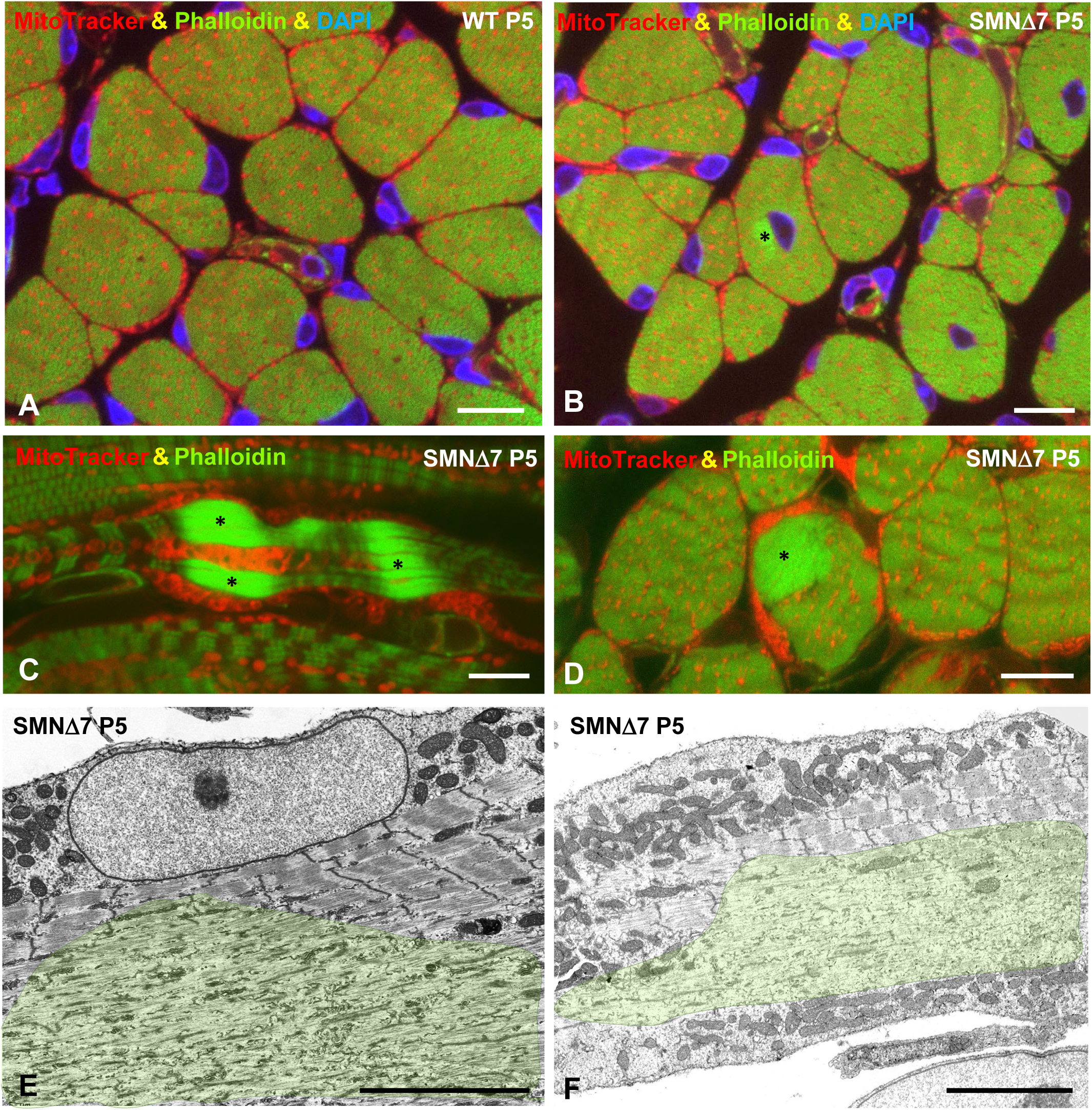
**(A-C)** Representative confocal images of transversal (A, B.D) and longitudinal (C) cryosections from TA myofibers double or triple stained to label mitochondria (Mitotracker, red channel), thin actin filaments (FITC-phalloidin, green channel) and nuclei (DAPI, blue channel) in WT (A) and SMNΔ7 (B-D) samples at PND (P5). **(A)** Typical WT myofibers with peripheral myonuclei and intermyofibrillar and peripheral mitochondria (red spots). **(B-D)** Presence of some central myonuclei (B) and mitochondria-free bright green fluorescent foci of F-actin accumulations (asterisks) in SMNΔ7 myofibers. **(E-F)** Electron microscopy images of myofibers from SMNΔ7 mice at PND stage (P5) showing extensive sarcoplasmic areas of disarrayed myofilaments and unrecognized sarcomeres free of mitochondria (transparent green areas). Scale bar: 10 μm (A, B, C, D) and 5 μm (E, F).

## DISCUSSION

The classical paradigm in SMA myopathy is that MN loss and muscle denervation have major effects on muscle atrophy and motor dysfunction (Lunn and Wang, 2008; Prior, 2010). However, increasing evidence in cellular and animal models of SMA supports that SMN deficiency also induces cell-autonomous alterations in the maturation and function of myoblasts/myotubes, myofibers, and muscle satellite cells (Boyer et al., 2013a; Bricceno et al., 2014; Cifuentes-Diaz et al., 2001; Hayhurst et al., 2012; Hellbach et al., 2018; Martínez-Hernández et al., 2009; Mutsaers et al., 2011; Shafey et al., 2005).

To better understand SMA myopathy, it is essential to explore in depth the cellular and molecular changes occurring in muscle at early stages of disease, prior to muscle denervation and MN loss (PND stage). Most of the previous studies in SMN-deficient muscle have focused on changes in molecular phenotype, including *i)* defects in the myogenic program (Boyer et al., 2014; Bricceno et al., 2014; Hayhurst et al., 2012; Hellbach et al., 2018; Martínez-Hernández et al., 2009); *ii)* alterations in the molecular organization of the cytoskeleton, contractile machinery and mitochondria (Bricceno et al., 2014; Chemello et al., 2023; Motyl et al., 2020; Mutsaers et al., 2011); *iii)* sarcolemmal and DNA damage (Cifuentes-Diaz et al., 2001; Mutsaers et al., 2011); *iv)* changes in the proteomic profile (Boyer et al., 2013b; Motyl et al., 2020); and *v)* dysfunction of neuromuscular synaptic transmission (Dachs et al., 2011; Franco-Espin et al., 2022; Kong et al., 2009; Lee et al., 2011; Murray et al., 2008). Additionally, SMN expression modifiers were identified using genome-wide RNAi screening (McCormack et al., 2021). In this study, we used the high resolution provided by electron microscopy to characterize for the first time the ultrastructural alterations in sarcomeres (contractile machinery) and SR and triads (excitation–contraction coupling machinery) underlying early postnatal nonatrophic myopathy in the SMNΔ7 mouse. We also provided new insights into the changes occurring in the phenotype and fusion dynamics of SMA-myofiber mitochondria and characterized the expression levels of several genes potentially involved in this nonatrophic myopathy. We report these changes at P5, a representative time point of the PND stage in SMNΔ7 mice (Boyer et al., 2013b; Khayrullina et al., 2020). Our results indicate that SMN deficiency causes early ultrastructural alterations in myofibers, which are independent of muscle denervation and MN loss. These alterations may result in the dysfunction of myofiber contractile properties and contribute to early onset motor defects in the SMA (see Graphical Summary).

### SMN deficiency leads to nonatrophic myopathy in SMN**Δ**7 mice at the PND stage

An important finding of our study was the absence of myofiber atrophy in the TA muscle of SMNΔ7 mice at the PND stage. This was evidenced by the lack of significant differences in myofiber size between SMNΔ7 and WT samples at P5. Furthermore, as several studies indicate that muscle atrophy reflects an imbalance between anabolic and catabolic pathways (Masiero et al., 2009; Wing et al., 2011), we analyzed the expression levels of genes encoding proteins involved in both pathways. In support of the early nonatrophic SMA myopathy evidenced by morphometric studies, we found that the mRNA expression level of *IL-15* did not change in the SMNΔ7 TA muscle compared to that in WT samples during the PND stage. The strong anabolic effect of IL-15 on muscle cells is supported by the demonstration that its overexpression in differentiating myotubes induces 5-fold greater levels of sarcomeric α-actin (Quinn et al., 2002). In contrast, nonsignificant changes in the expression levels of the atrogenes *MuRF1* and *Atrogin-1*, which encode two ubiquitin ligases in the UPS that are implicated in myofilament protein proteolysis, were observed in TA muscle samples from SMNΔ7 and WT mice (Bodine et al., 2001; Cohen et al., 2009; Sacheck et al., 2004). We hypothesize that the increased expression of the promyogenic gene *Dcn,* which negatively regulates *Atrogin-1* mRNA levels (Kanzleiter et al., 2014; Marshall et al., 2008), prevents the upregulation of the atrogin-1 E3 ligase and the subsequent ubiquitination of its sarcomeric protein targets. Thus, unlike the upregulation of atrogenes associated with skeletal muscle atrophy (Bodine et al., 2001), the lack of atrogene-dependent proteolysis during the PND stage in SMNΔ7 mice could preserve myofiber size and thus contribute to nonatrophic early myopathy in SMA. Additionally, no ultrastructural signs of activation of the autophagyLlysosome system, a proteolytic pathway involved in muscle atrophy (Masiero et al., 2009), were observed in SMNΔ7 mice at the PND stage.

To further investigate the potential contribution of changes in the myogenic program to the absence of muscle atrophy, we evaluated the expression of the main myogenesis regulator factors (MRFs) *Pax7*, *MyoD, Myog* and *Mrf4* in the TA muscle from SMNΔ7 mice at P5. Previous studies in purified myoblasts and muscle from SMA mouse models and patients concluded that MRF expression is perturbed when SMN levels are reduced (Boyer et al., 2014; Bricceno et al., 2014; Hayhurst et al., 2012; Ripolone et al., 2015). However, differential changes in MRF expression patterns have been found in these studies (Jha et al., 2023). For instance, whereas Boyer et al. (2014) reported decreased Pax7, MyoD and myogenin levels in muscles from SMA mice during the symptomatic period, Bricceno et al. (2014), using transformed myoblasts derived from the same SMA mouse line, demonstrated that decreased Pax7 expression was accompanied by increased MyoD and myogenin levels. The upregulation of *Pax7,* a transcription factor involved in regenerative myogenesis (Collins et al., 2009; Seale et al., 2000), and the presence of mitotic satellite cells in SMA muscles found in our study are consistent with a potential regenerative response to myofiber lesions. Moreover, the unchanged expression of *MyoD, Myog* and *Mrf4* in the TA muscle of SMNΔ7 mice compared to that in age-matched WT samples suggested that the myogenic program was still active during the PND stage. Similarly, Ripolone et al. (2015) reported increased levels of three MRFs (MYF5, MYOD, and MYOG) in the muscle of patients with SMA. In conclusion, the lack of activation of two key catalytic pathways, atrogene-dependent UPS proteolysis and the autophagy-lysosomal system, together with the sustained activity of the myogenic program, are congruent with early nonatrophic myopathy in SMNΔ7 mice during the PND stage.

### Phenotypic ultrastructural alterations of the contractile machinery underlying motor disturbances in SMN**Δ**7 mice: sarcomere disruption associated with actinopathy

The major ultrastructural alterations found in the SMA myofibers of SMNΔ7 mice consisted of both focal and segmental disruptions of sarcomere architecture. The time course of sarcomeric lesions appears to be as follows: *i)* misalignment of myofibrils, *ii)* disassembly of Z-discs, *iii)* disarrangement of myofilaments with disappearance of the typical banding pattern in sarcomeres, and *iv)* abnormal accumulation of F-actin. These early myofiber lesions, already observed at P1, could act as physical obstacles to maximal contraction force generation and, therefore, impede skeletal myofiber output. Additionally, it is worth noting the presence of myofiber segments with overcontracted minisarcomere repeats, where no force is generated due to the excessive overlap of actin and myosin filaments that impedes them from sliding over each other (Duchateau and Enoka, 2016). These structural alterations of sarcomeres observed in SMNΔ7 mice have also been reported to occur in human SMA myofibers (Berciano et al., 2020a) and could be considered a pathological structural hallmark of SMA myopathy. Indeed, previous studies have reported the localization of SMN in mouse and human sarcomeres, suggesting that the SMN protein is involved in the maintenance of sarcomere architecture (Berciano et al., 2020a; Rajendra et al., 2007; Walker et al., 2008).

The disorganization of myofilaments and aberrant accumulation of F-actin in myofibers are both ultrastructural features of myofibrillar myopathies and are characterized by altered F-actin expression, polymerization and dynamics, resulting in the aberrant assembly of thin myofilaments (Batonnet-Pichon et al., 2017; Dubowitz and Sewry, 2007b; Nowak et al., 2013). The pathological accumulation of actin thin filaments has also been reported in other myopathies and muscle disorders, such as progressive actin-accumulation myopathy caused by mutations in the *Actin Alpha 1, Skeletal Muscle (ACTA1*) gene (Bornemann et al., 1996; Goebel et al., 1997; North and Laing, 2008; Nowak et al., 2013). We propose that the early postnatal myopathy in SMNΔ7 mice with abnormal accumulations of F-actin could be considered SMN deficiency-induced actinopathy. Indeed, previous molecular studies in cellular and murine models of SMA have reported the disruption of skeletal muscle actin-cytoskeleton signaling pathways, the upregulation of the *Acta1* gene, and the dysregulation of the RhoA pathway (Motyl et al., 2020; Mutsaers et al., 2011). In this regard, the RhoA/ROCK pathway is a key regulator of F-actin assembly and dynamics through the phosphorylation of downstream F-actin targets, including myosin light chain phosphatase, cofilin and profilin (Da Silva et al., 2003; Grandy et al., 2022; Nölle et al., 2011). Importantly, several studies support that dysregulation of the ROCK pathway contributes to SMA pathogenesis through direct interaction of SMN with profilin and regulation of its activity during F-actin polymerization (Bowerman et al., 2010; Hensel and Claus, 2018; Nölle et al., 2011). Indeed, ROCK inhibitors have been proposed for SMA therapy (Bowerman et al., 2010; Coque et al., 2014).

### Potential influence of myonuclear domains on the nonrandom distribution of myofiber lesions in SMN**Δ**7 mice

The cellular mechanisms that determine the distribution of lesions along SMNΔ7 myofibers are unknown. We propose that the spatial localization of lesions may correspond to sarcoplasmic regions under the specific influence of one or several dysfunctional “myonuclear domains”. The polyploid and multinucleate nature of skeletal myofibers has led to the concept of a “myonuclear domain”, which states that each myonucleus regulates a discrete volume of sarcoplasm, for example, by supplying ribosomal particles and mRNA transcripts. This principle also provides a logical explanation for establishing internuclear distances and is based on the notion that DNA content and cell volume are tightly coupled to maintain the proper nucleus–cytoplasm ratio (Gregory, 2001). Insights from modern technologies, including muscle spatial transcriptomics, single-myonuclei RNA sequencing and fluorescently tagged proteins, have provided further information on the RNA and protein mobility restrictions in muscle cells and their relationship with myonuclear domains (Kann and Krauss, 2019; McKellar et al., 2021). For instance, Morin et al. (2023) demonstrated, by means of fluorescent protein tagging, that dystrophin is highly compartmentalized to myonuclear-defined sarcolemmal domains. Similarly, another study reported that mRNA transcripts of *AChR* subunit genes exhibit restricted mobility within specialized synaptic myonuclear domains, which refer to those myonuclei located beneath the postsynaptic sarcolemma (Merlie and Sanes, 1985). The assumption of dysfunction of myonuclear domains is consistent with our observation of sarcoplasmic areas where both disorganized and normal sarcomeres coexist, with a sharp transition between them. We hypothesize that the existence of myonuclear domains is more prone to injury and, potentially, more vulnerable to low SMN levels. However, further studies using innovative technologies are needed to determine the molecular basis of these distinct myonuclear domains and their differential contributions to the pathogenesis of SMA myopathy.

### Changes in the ultrastructural phenotype of the excitation-contraction coupling machinery in SMN**Δ**7 mice

To our knowledge, the present study evaluated for the first time the ultrastructural alterations in the excitation-contraction coupling machinery (SR and triads) of myofibers at early stages of SMA myopathy. Briefly, the SR plays a key role in Ca^2+^ homeostasis by removing Ca^2+^ from the cytosol. This process is carried out by Ca^2+^ ATPases (SERCAs), which pump Ca^2+^ from the cytosol to the SR lumen. Thus, the SR stores, releases, and reuptakes Ca^2+^ to sustain an effective contraction-relaxation cycle. Two terminal cisterns of the SR are assembled with a T-tube to form the “triad”, the structural platform responsible for transducing the depolarization of the sarcolemma into Ca^2+^ release from the SR to trigger muscle contraction. This mechanism is known as excitationLcontraction coupling (for review, see (Dulhunty, 2006; Rossi et al., 2022).

Dilation of the terminal cisterns of triads with preserved T-tubules and moderate swelling of the longitudinal network of the SR are the earliest ultrastructural changes we found in SMNΔ7 mouse myofibers prior to sarcomere disruption. These phenotypic alterations in the excitation-contraction coupling machinery suggest Ca^2+^ homeostasis perturbation, which could contribute to the motor disturbances and muscle weakness found in the early stages of disease (PND stage) (Supplementary Video 1, 2). Moreover, we occasionally observed signs of vacuolar degeneration of the SR but not of mitochondria, which is consistent with osmotic perturbation presumably caused by a severe ion imbalance between the lumen of the SR and the cytosol. In myofiber areas with sarcomere disorganization, the network of SR cisterns appeared dysmorphic and fragmented, and the triads were disassembled or absent. These findings clearly reflect a massive impairment of the excitation-contraction coupling function linked to the disorganization of sarcomere architecture.

Dysregulation of Ca^2+^ homeostasis, accompanied by a decrease in the rate of Ca^2+^ uptake and an increase in the concentration of cytosolic Ca^2+^, has been associated with SR dilation in experimental models of exercise-induced muscle damage (Byrd, 1992). Similar findings have been observed in muscular disorders, such as malignant hyperthermia and central core disease caused by mutations in genes encoding the following: *i)* Ca^2+^ release channels (the ryanodine receptors, *RYR*), *ii)* calsequestrin (*CASQ*) and *iii)* the dihydropyridine receptor (*CACNA1s*), a voltage-dependent L-type Ca^2+^ channel located on the T-tubule (MacLennan and Zvaritch, 2011). In SMA mice and SMN-deficient cell lines, alterations in the molecular phenotype of the excitation–contraction coupling machinery have been shown in cardiomyocytes and skeletal myofibers. In cardiomyocytes from SMNΔ7 mice and SMA patient-derived iPSCs, (Khayrullina et al., 2020)) reported a reduced expression of SERCA2, which resulted in the blockade of Ca^2+^ reuptake in the SR and led to an impairment of cardiomyocyte function. Similarly, in the TA muscle of *Smn*^-/-^;*SMN2* mice, Boyer et al. (2013b) reported a decrease in the level of SERCA1a, the predominant SERCA isoform found in the fast-twitch TA muscle (Beard et al., 2004). Moreover, the authors reported dysregulated expression of spliced RyR1 isoforms, which probably led to reduced Ca^2+^ release from the SR (Boyer et al., 2013b). In conclusion, the alterations in the ultrastructural phenotype of the excitation-contraction coupling machinery reported here provide significant insights into Ca^2+^ dysregulation as an important contributor to SMA pathophysiology in the early stages of this disease, prior to muscle denervation and MN loss.

### Upregulation of mitochondrial biogenesis and fusion dynamics in SMN**Δ**7 mice at the PND

Our findings in the TA muscle of SMNΔ7 mice at the PND stage of the disease strongly support the potential role of SMN in the maintenance of the mitochondrial phenotype. Different patterns of mitochondrial behavior have been reported in the SMA. Several studies in animal and cell culture models of this disease have shown a deficit of oxidative phosphorylation accompanied by impaired complex I and IV activity, increased mitochondrial ROS production, mtDNA depletion, and structural damage to mitochondria (Berger et al., 2003; Chemello et al., 2023; James et al., 2021; Ripolone et al., 2015; Zilio et al., 2022). Here, we found that the mitochondrial phenotype of SMA myofibers is determined by the activation of mitochondrial biogenesis and fusion dynamics, preservation of structural integrity, and lack of mitophagy. The divergences between the mitochondrial patterns described by us and others in SMA myofibers could result from the different developmental periods examined (PND and ND stages).

The mitochondrial phenotype we observed in SMNΔ7 TA myofibers during the PND stage consisted of an enlargement of the mitochondrial compartment and the presence of very long intermyofibrillar mitochondria. This, in addition to the presence of interacting mitochondria, suggests the activation of both dynamic mitochondrial biogenesis and fusion processes. Accordingly, key genes involved in these processes, such as *Mfn1*, *Mfn2,* and *PGC-1a*, were upregulated in the SMNΔ7 TA muscle at the PND stage. The upregulation of *PGC-1a*, a master gene of mitochondrial biogenesis (Halling and Pilegaard, 2020; Romanello et al., 2010), has been reported to induce *Mfn2* transcription and translation through its interaction with *Mfn2* (Emery and Ortiz, 2021; Soriano et al., 2006). Furthermore, we found increased expression of *Fdcn5* (irisin), a molecular component of the PGC-1α/FDCN5/irisin signaling pathway, which is also involved in mitochondrial biogenesis and function (Chen et al., 2017; Srinivasa et al., 2016; Vaughan et al., 2015). Collectively, these findings suggest that, during the PND stage, SMN deficiency overactivates mitochondrial biogenesis and fusion in myofibers of the TA muscle, which, under physiological conditions, is a predominantly fast-twitch glycolytic muscle (mainly composed of type IIX and type IIB fibers) that displays low fusion mitochondrial rates (Hamalainen and Pette, 1993; Mishra and Chan, 2016).

We propose that the upregulation of *Mfn1*, *Mfn2,* and *PGC-1a* found in our study could promote the oxidative capacity of TA myofibers, which is a compensatory mechanism that sustains bioenergetic metabolism under conditions of motor dysfunction caused by focal and segmental myofiber lesions. Indeed, ultrastructural signs suggestive of mitochondrial dysfunction, such as matrix swelling, cristae disruption, vacuolization, and the formation of mitochondria-derived vesicles (He et al., 2020; Sugiura et al., 2014), were occasionally found in the mitochondria of SMNΔ7 myofibers at the PND stage. The only ultrastructural feature potentially related to mitochondrial dysfunction was the interaction between intact mitochondria and swollen SR cisterns (Figs. 6D and S2), which could impair the normal flux of Ca^2+^ from the SR to mitochondria required for ATP production regulation (Rossi et al., 2022). The normal interaction of mitochondria with the SR is mediated by tethering structures that link the mitochondrial outer membrane to the SR, as well as by the voltage-dependent anion channel 1 and RyR (Boncompagni et al., 2009; Min et al., 2012). Briefly, RyR opening increases the Ca^2+^ concentration in local microdomains, allowing the transfer of Ca^2+^ to the mitochondrial matrix (Mammucari et al., 2018). This mechanism could be impaired by SR swelling, resulting in defective bioenergetic signaling. Notably, our results revealed the absence of ultrastructural signs of mitophagy, a cellular mechanism usually coupled with the activation of mitochondrial fission dynamics (Song et al., 2015). Finally, mitochondria are commonly absent in myofiber lesions. We think that the local disruption of myofibrils and their intermyofibrillar spaces containing bundles of longitudinally oriented microtubules (Denes et al., 2021) could prevent the microtubule-based transport of intermyofibrillar mitochondria, resulting in local depletion of mitochondria and a reduction in ATP bioenergetics.

The elevated rates of mitochondrial biogenesis and fusion found in the TA myofibers of SMNΔ7 mice, suggestive of an enhancement of oxidative metabolism, do not correlate with TA fiber type switching toward higher levels of slow MyHC-I fibers. Conversely, the SMNΔ7 TA muscle displayed the same low number of MyHC-I fibers as did the TA muscle in age-matched WT animals. Consistent with this view, previous studies have shown that enhanced mitochondrial fusion ex vivo is not accompanied by fiber-type switching and that mitofusin deletion does not prevent the formation of oxidative fibers (Chen et al., 2010; Mishra and Chan, 2016; Pette and Staron, 2000). Thus, we propose that alterations in mitochondrial morphology and energy metabolism occurring in SMN-deficient myofibers at the PND stage do not play a direct role in determining the fiber type and do not appear to be synchronized with the rearrangement of the myofiber myosin profiles.

Overall, our study provides new data for a better understanding of the cellular basis underlying skeletal SMA myopathy induced by low SMN levels. Dysfunction of myofibers during early nonatrophic myopathy, prior to muscle denervation and MN loss (PND stage), could retrogradely influence the pathophysiology of secondary atrophic (neurogenic) myopathy accompanied by muscle denervation (ND stage). The present results also indicate that skeletal muscle is a major therapeutic target for improving motor function in SMA in combination with therapies aimed at restoring SMN levels in MNs.

## ACKNOWLEDGEMENTS

The authors wish to thank Raquel García-Ceballos, Lídia Piedrafita, Sílvia Gras and Kimi Bencosme for technical assistance.

## Ethical guidelines statement

All animal studies have been approved by the appropriate ethics committee and have therefore been performed in accordance with the ethical standards laid down in the 1964 Declaration of Helsinki and its later amendments.

---

**Supplementary Figure 1. (A)** Western blotting analysis of SMN expression in TA muscle lysates from WT and SMNΔ7 mice at the indicated postnatal ages (P). Lamin A/C was used as load control. Protein expression fold of SMN protein is indicated. **(B)** Quantitative analysis of the mean number of αMNs present in transversal cryosections of spinal cords stained with propidium iodide from WT (n=6) and SMNΔ7 (n=6) mice at the indicated postnatal ages (P). *p* values from WT and SMNΔ7 data comparison were 0.0865, 0.0727, 2.6-04 and 9.8E-11 at P0, P5, P10 and P14, respectively. **(C)** Quantitative analysis of body weight changes in WT (n=12) and SMNΔ7 (n=12) mice at indicated postnatal ages. **(D)** Representative confocal images of transversal cryosections of anterior horns stained with propidium iodide from WT and SMNΔ7 mice at P5 and P14 used for the αMN quantification shown in panel B. Schematic illustration of the time-course of pre-neurodegenerative (PND) and neurodegenerative (ND) stages. Scale bar = 200 μm in (D).

**Supplementary Figure 2. (A)** Electron micrograph showing the ultrastructural phenotype and organization of subsarcolemmal and intermyofibrillar mitochondria in a WT myofiber at P5. Z: Z-discs. **(B-D)** Fine structure of mitochondria in SMNΔ7 myofibers. **(B)** Clusters of intermyofibrillar mitochondria illustrating the direct link between the outer membranes of adjacent mitochondria (arrows) and the interactome of mitochondria with both lipid droplets (LD) and dilated SR cisterns (asterisk). **(C)** High magnification detail of the close interaction between the outer mitochondrial membrane and the membrane of a dilated SR cistern (asterisk). Note the well-preserved ultrastructure of mitochondria. **(D)** Great accumulation of subsarcolemmal mitochondria of variable size and morphology. Note the absence of mitophagy images. LD: lipid droplets. Scale bar = 2 μm (A, B), 200nm (C) and 5 μm (D).

