## Supplementary figures and images for "SMN deficiency induces an early non-atrophic myopathy with alterations in the contractile and excitatory coupling machinery of skeletal myofibers in the SMNΔ7 mouse model of spinal muscular atrophy"

### Supplemental Figures 1 and 2

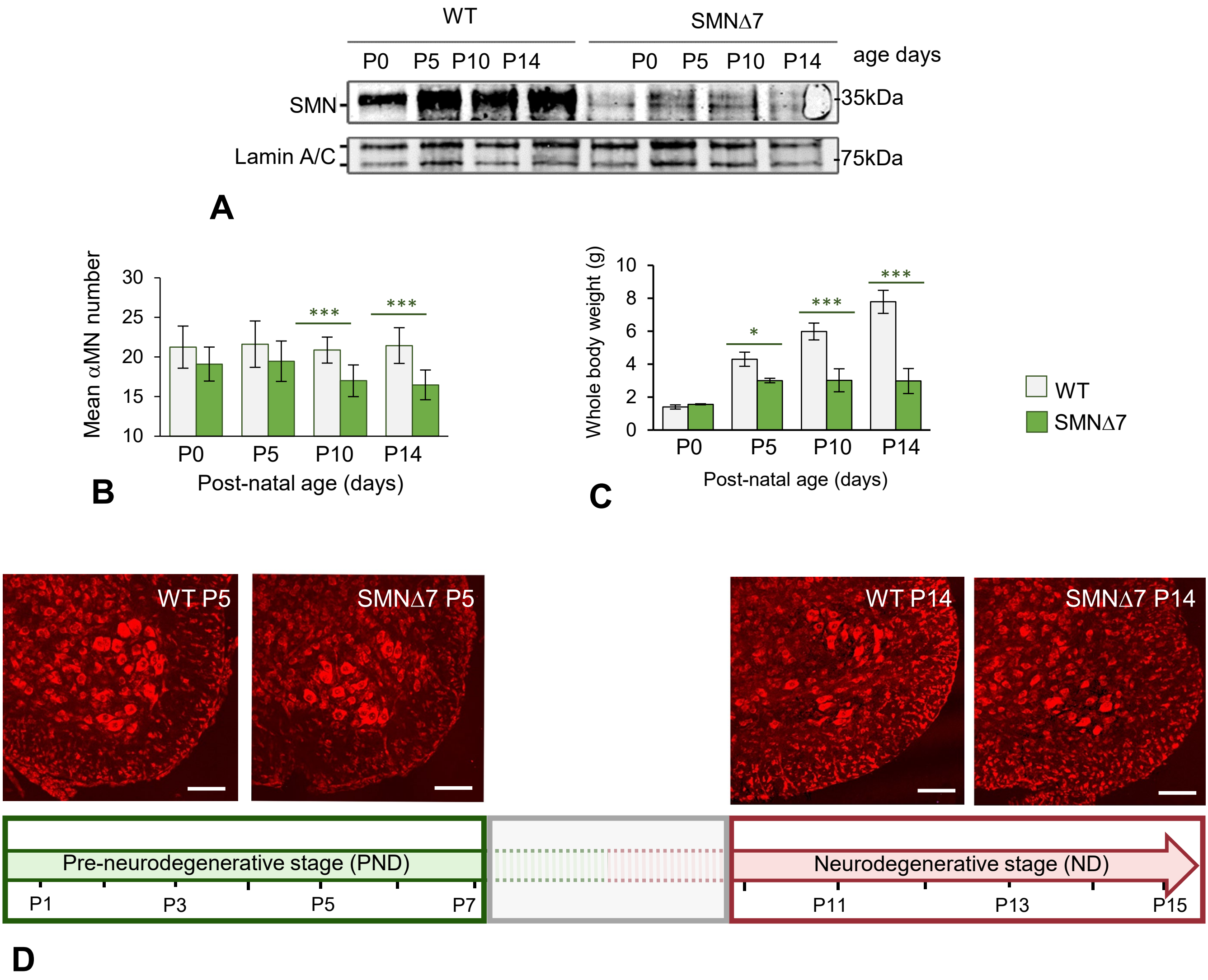


Supplementary Figure 1

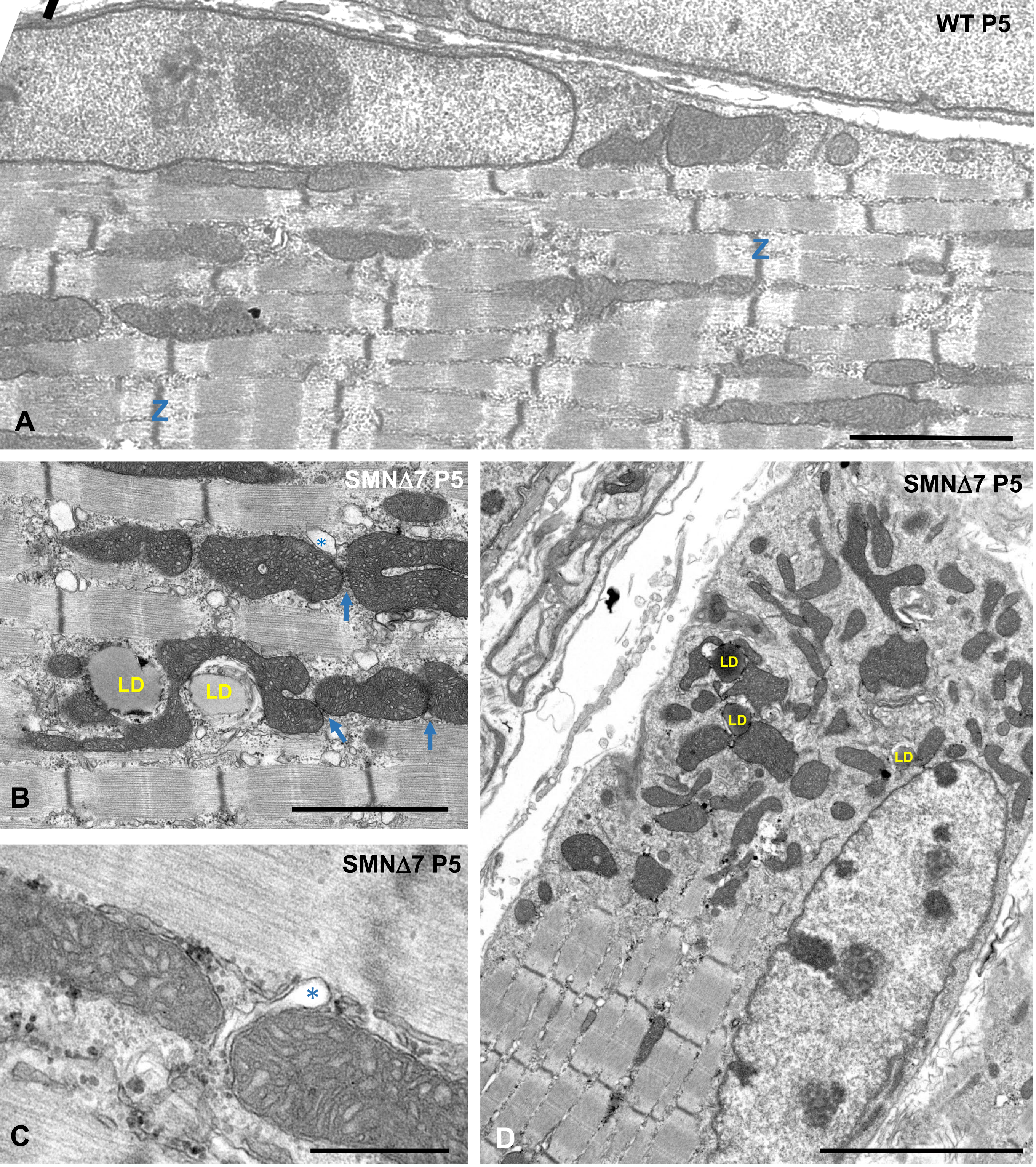


Suplementary Figure 2
